# CalTrig: A GUI-based Machine Learning Approach for Decoding Neuronal Calcium Transients in Freely Moving Rodents

**DOI:** 10.1101/2024.09.30.615860

**Authors:** Michal A. Lange, Yingying Chen, Haoying Fu, Amith Korada, Changyong Guo, Yao-Ying Ma

**Author notes:** **Corresponding authors**: Dr. Yao-Ying Ma & Dr. Changyong Guo, Department of Pharmacology and Toxicology, Indiana University School of Medicine 635 Barnhill Drive, Indianapolis, IN 46202.

## Abstract

Advances in *in vivo* Ca^2+^ imaging using miniature microscopes have enabled researchers to study single-neuron activity in freely-moving animals. Tools such as MiniAN and CalmAn have been developed to convert **Ca**^2+^ **v**isual signals **to n**umerical data, collectively referred to as CalV2N. However, substantial challenges remain in analyzing the large datasets generated by CalV2N, particularly in integrating data streams, evaluating CalV2N output quality, and reliably and efficiently identifying Ca^2+^ transients. In this study, we introduce CalTrig, an open-source graphical user interface (GUI) tool designed to address these challenges at the post-CalV2N stage of data processing. CalTrig integrates multiple data streams, including Ca^2+^ imaging, neuronal footprints, Ca^2+^ traces, and behavioral tracking, and offers capabilities for evaluating the quality of CalV2N outputs. It enables synchronized visualization and efficient Ca^2+^ transient identification. We evaluated four machine learning models (i.e., GRU, LSTM, Transformer, and Local Transformer) for Ca^2+^ transient detection. Our results indicate that the GRU model offers the highest predictability and computational efficiency, achieving stable performance across training sessions, different animals and even among different brain regions. The integration of manual, parameter-based, and machine learning-based detection methods in CalTrig provides flexibility and accuracy for various research applications. The user-friendly interface and low computing demands of CalTrig make it accessible to neuroscientists without programming expertise. We further conclude that CalTrig enables deeper exploration of brain function, supports hypothesis generation about neuronal mechanisms, and opens new avenues for understanding neurological disorders and developing treatments.

## 1. INTRODUCTION

Neurons within the same brain region are diverse in type, connectivity, and activity, responding to stimuli with high temporal precision. One of the key advances in modern neuroscience is the shift from studying brain regions as functional units to focusing on individual neurons. Monitoring single-neuron activity in freely-moving animals allows for deeper insights into brain function and dysfunction. Recent advances in *in* vivo imaging and fluorescent Ca^2+^ indicators, coupled with miniature microscopes (miniScopes), have revolutionized the study of neural dynamics in freely-moving animals. Tools have been developed to convert Ca^2+^ visual signals (i.e., Ca^2+^ imaging video files) to numerical data (e.g., changes in fluorescent signal intensity indicating the Ca^2+^ influx for each neuron), denoted **CalV2N**. Early tools such as Suite2P^1^, SIMA ^2^, STNeuronNet ^3^, CalmAn ^4^, were created to process two-photon imaging data. Due to the low resolution and poor signal-to-noise ratio (SNR) in one-photon imaging data, more specific CalV2N tools were created including MIN1PIPE^5^ and MiniAn^6^. The algorithms used in CalV2N tools started with principal-component analysis/independent component analysis (PCA/ICA)^7^, and then upgraded to constrained non-negative matrix factorization (CNMF), and its various derivations such as CNMF-E^6^. Due to better reliability in demixing the activities of overlapping cells, the computational efficiency and accessibility of parameter adjustment, CNMF based MiniAn has become a reliable choice and selected as the CalV2N tool to extract Ca^2+^ traces in this study.

Despite the availability of CalV2N tools, substantial challenges remain in drawing meaningful conclusions from Ca^2+^ imaging data. **First**, the lack of synchronized visualization of multiple data streams: Usually at least two original data streams are collected, including behavioral video tracking and the Ca^2+^ image. After data processing by the CalV2N tool, three data streams are generated, including extracted Ca^2+^ traces, footprint of identified cells, and the processed video. The current available CaV2N tools are usually Python-based. Visualization of different lines of data or videos are primarily contained in discrete sections (e.g., background removal, seed selection, and other sections in MiniAn serving as an example CalV2N tool). Although this provides insight into the effects of single parameters, an integrative visual platform is missing that would provide the user with a review of the multiple lines of data. **Second**, the global parameters applied across the data often result in inconsistencies, as signal quality varies between neurons. The parameters that are available to the user are applied across the entirety of the input data, which can contribute to mixed results to the quality of extracted Ca^2+^ traces. MiniAn for instance, visualizes 5-10 cells at random to ascertain the impact of a parameter. In our research group, we observed inconsistencies arising from differing SNRs across different cells and varying intensities of resultant Ca^2+^ traces, which has created a demand for a post-CalV2N tool. **Third**, the verification of detected cells and extracted Ca^2+^ traces remains uncertain. **Forth**, there is no well-established tool available to time efficiently and reliably identify Ca^2+^ transients. **Fifth**, we would also like to highlight that the challenges mentioned are significantly amplified by the complexity and sheer volume of data processed through CalV2N. This includes: (1) high temporal resolution, with sampling rates of 10-60 Hz; (2) high spatial resolution, with pixel sizes ranging from 0.8-1.0 µm and a field of view (FOV) up to 1.0 mm × 0.8 mm; (3) high cell throughput, with 50-200 neurons typically detected per animal, resulting in massive datasets, especially when scaled across multiple animals (e.g., 10-20 per group) and various experimental conditions; and (4) integration with behavioral data, adding further complexity to the analysis.

To address these challenges at the post-CalV2N stage, we developed the Ca^2+^ transient identifier GUI (CalTrig), a graphical user interface (GUI) using the Python package PyQT5. CalTrig integrates all outputs from MiniAn (including Ca^2+^ imaging original processed videos, Ca^2+^ traces, cell footprints, and behavioral videos) to facilitate synchronized visualization, evaluate the performance of CalV2N, and identify Ca^2+^ transients. The objective of creating CalTrig is listed below:

- **Efficient performance**: Low computing demands ensure smooth operation on standard computers.
- **User-friendly interface**: Neuroscientists without programming skills can explore and analyze data at neuronal levels with high temporal resolution.
- **Integrative visual exploration**: CalTrig enables evaluation of CalV2N performance and reliable detection of Ca^2+^ transients by integrating multiple data streams and videos.
- **Multiple options for Ca^2+^ transient detection**: CalTrig supports (1) manual, (2) parameter-based, and (3) machine learning-based detection methods.
- **Integration of detection strategies**: Detection strategies can be combined. For example, parameter-based autodetection or machine learning outputs can be manually corrected, and parameter-based autodetection can serve as a preliminary step before manual detection, helping set up datasets for training machine learning models.
- **Informative output**: Detailed Ca^2+^ transient data for each neuron and an overview of a pool of neurons from one mouse brain can be exported as data tables or figures for publication.

In summary, CalTrig is developed to bridge the gap between CalV2N tools and the final stages of data analysis, simplifying the workflow and enabling researchers to extract meaningful biological insights from raw data. Although the potential readership is broad, this research article primarily aims to assist neurobiologists in processing *in vivo* Ca^2+^ imaging data collected using a single-photon miniScope. We would like to give a brief introduction to the article’s organization, which differs from a typical neurobiological research paper. In the **Introduction**, we elaborate on the significance of *in vivo* Ca^2+^ imaging in freely-moving animals and discuss the challenges encountered after extracting Ca^2+^ traces using tools like CalV2N and MiniAn. We then conclude stating the goals for developing CalTrig. The **Methods** section not only reiterates previously established procedures for collecting Ca^2+^ imaging data from freely-moving mice (with more details available in the **Supplementary Documents**) and extracting the Ca^2+^ traces, as we reported earlier^8,9^, but also provides detailed information about the newly developed tool, CalTrig. The latter includes data loading from CalV2N, data visualization, cell verification, CalV2N evaluation, Ca^2+^ transient identification, and exporting figures or data for statistical analysis, ready for publication. We present three strategies for Ca^2+^ transient identification (i.e., parameter-based, manual, and machine learning-based detection), including their procedures and applications. In the **Results** section, we compare the performance of multiple machine learning models, evaluate whether the established machine learning model can be used to identify transients across datasets collected at different time points, from different animals, and in various brain regions. Finally, the **Discussion** highlights the advantages of using machine learning models, the integrative visual exploration interface, and the unique features of CalTrig.

## 2. METHODS

### 2.1. *In vivo* Ca^2+^ Imaging data collection

#### 2.1.1. Experimental animals

All *in vivo* procedures on laboratory animals were performed in accordance with the United States Public Health Service Guide for Care and Use of Laboratory Animals and were approved by the Institutional Animal Care and Use Committee at Indiana University School of Medicine. Ten male C57BL/6J, bred in-house using breeders originally derived from the Jackson Laboratory, were used in this study. The mice were group housed, except for those with GRIN lens implants, which were singly housed to prevent cage mates from damaging the implanted lens. All mice had free access to chow and water in home cage, maintained on a 12-hour light/dark cycle (light on at 7:00 AM and off at 7:00 PM).

#### 2.1.2. Surgical procedures: Microinjection of AAV

Mice were anesthetized with 2.5% isoflurane for induction and maintained with ∼1.2%. A 28-gauge injection needle was used to unilaterally inject the tAAV1-Syn-jGCaMP8f-WPRE or AAV1-CaMKIIa-jGCaMP8f-WPRE solution (0.5 µl/site, 0.1 µl/min) *via* a Hamilton syringe into the secondary motor cortex (M2) (coordinates in mm: AP, +1.80; ML, ±0.60; DV, −1.30) or the medial prefrontal cortex (mPFC) (coordinates in mm: AP, +2.05; ML, ±0.3; DV, −2.45) using a Pump 11 Elite Syringe Pumps (Harvard Apparatus). Injection needles were left in place for 5 min following injection.

##### Lens implantation

A couple of minutes after withdrawing the AAV injection needle, a unilateral GRIN lens (Inscopix Inc, #1050-004595, Diameter: 1.0 mm; length: ∼4.0 mm; Working Distance: 200 µm) was lowered through the cranial window to 200 µm above the center of the virus injection site. The open space between the lens’ side and the skull opening was sealed with surgical silicone (Kwik-Sil) and secured by dental cement (C&B Metabond). The exposed part of the lens above the skull was further coated with black cement (Lang Dental Mfg. Co.’ Inc.).

##### Base-plating

3 weeks later, mice were anesthetized with isoflurane again. The cement on top of the GRIN lens was carefully removed using drill bits until the lens was exposed. The top of the lens was then cleaned using lens paper and a cleaning solution. A metal baseplate was mounted onto the skull over the lens using Loctite super glue gel, guided by a MiniScope V4.4 (Open Ephys) for optimal field of view. Once the baseplate was securely mounted, the MiniScope was removed. A protective cap was attached to the baseplate, and mice were returned to home cage.

##### Verification of AAV expression and the lens location

After the completion of the *in vivo* Ca^2+^ imaging, M2 or mPFC-containing coronal slices were prepared as described before^10^, then fixed in 4% Paraformaldehyde (PFA) for no less than a couple of hours. After a brief rinse with PBS, slices were mounted with Prolong^TM^ Gold antifade mounting reagent with DAPI (Invitrogen, Cat# P36931). Confocal imaging was performed using a Zeiss LSM 800 confocal microscope. The criteria for animal inclusion in this study were (a) highly enriched AAV expression in mostly pyramidal neurons, within-M2 or mPFC viral injection site, and (b) the footprint of the GRIN lens tip at the top of the targeting brain area.

#### 2.1.3. *In vivo* Ca^2+^ imaging recording

Mice were habituated to the *in vivo* Ca^2+^ recording procedure by mounting the miniScope V4.4 (Open Ephys) to the pre-anchored baseplate and recording for 5 min per day at the home cage for 3 days before starting the 1-hr daily recordings in an operant chamber (Med Associates). Data Acquisition (DAQ) box, supported by an open source, C^++^ and Open Computer Vision (OpenCV) libraries-based software, was used to collect both Ca^2+^ and behavioral video streams simultaneously controlled by the operant chamber software, MED-PC (Med Associates) *via* a TTL adaptor. The sampling frequency was 30 Hz.

### 2.2. Extraction of Ca^2+^ transient traces from the raw videos

Among multiple types of computational tools established previously to extract Ca^2+^ transients from raw videos, a Python-based analysis pipeline, Minian^11^, was used in our data analyses due to its low memory requirement and user friendly parameter options. In brief, there were five steps in the pipeline. First, multiple raw videos were batch loaded and subjected to a *PREPROCESSING* stage, where sensor noise and background fluorescence from scattered light were removed. Second, rigid brain motions were corrected by *MOTION CORRECTION*. Third, the initial spatial and temporal matrices for later steps were generated by a seed-based approach, called *SEEDS INITIALIZATION*. Fourth, the spatial footprints of cells were further refined. Fifth, the temporal signals of cells were also refined. The last two steps, the *SPATIAL UPDATE*, and the *TEMPORAL UPDATE*, as the core computational components based on CNMF algorithm, were repeated at least one more time.

### 2.3. CalTrig (Ca^2+^ Transient identification GUI)

CalTrig is python-based open source code with GUI interface. It is available in our lab Github station (https://github.com/cci-MaLab/Calcium-Transient-Analysis). The repository contains documentation, demos, and a message/discussion board. The code, which is compatible with Python 3.10, uses several open-source libraries including Xarray, Numpy, Pandas, PyQt5 and Pyqtgraph.

Computer system specifications for CalTrig development and testing: The development and testing of CalTrig were conducted on a system with the following specifications: CPU, Ryzen 9 7900X; RAM, 64GB; GPU, Nvidia RTX 4080; Operating System, Windows 11.

#### 2.3.1. Data loading

The front page of CalTrig serves as a hub for loading data (**Fig. S1A**). Data can be loaded by directly specifying the file path, or indirectly by loading the pre-generated INI (Initialization) files or JSON (JavaScript Object Notation) files.

The INI file uses a flat format with itemized information to detail a single data set. It incorporates both basic experimental design details (e.g., Animal ID, day, session stage) and multiple temporally synchronized data lines, including (1) the original and CNMF processed Ca^2+^ image video, (2) extracted Ca^2+^ trace data (i.e., the CNMF output), (3) behavioral data (i.e., operant behavioral such as active lever press, ALP; inactive level press, ILP; and Reinforcement, RNF), and (4) behavioral video tracking. The variables included in each CNMF dataset are listed in **Table 1** and included in Parameter list in CalTrig interface (**Fig. S2**). The INI file can be created by following the format provided in demo INI files, ensuring that both the experimental design and data streams are properly synchronized for exploration and analysis.

**Table 1.** CNMF variable.

| CNMF variable | Definition |
| --- | --- |
| **C** | Temporal matrix of $\text{Ca}^{2+}$ dynamics |
| **S** | Temporal matrix of $\text{Ca}^{2+}$ transient rising slope |
| **A** | Spatial matrix of cell footprints |
| **B** | Spatial matrix of background footprints |
| **f** | Temporal vector of background dynamic. |
| **YrA** | Temporal matrix of residual cell traces. |
| **Y/....** | A collection of 3D matrices containing original and processed videos. |
| **$\Delta F/F$** | Temporal matrix of $\Delta F/F$ values for cells. This is the only variable that is not considered mandatory, as the MiniAn pipeline does not generate it. When absent, it will be automatically generated in the CalTrig using a CAIMAN implementation. |

The JSON file, in contrast, uses a hierarchical format, representing the experimental design. Using data from one of our addiction projects as an example, the JSON file can collect information from multiple INI files, outlining details such as treatments for each animal, Ca^2+^ imaging days, and session stages. The JSON file can be generated by saving data after loading multiple INI files, either for longitudinal recordings in a single animal or for a group of animals assigned to the same experimental group.

#### 2.3.2. Data visualization

As shown in **Fig. S1B**, each set of data can be visualized in the window by their cell footprint and are arranged in a grid related to the experiment design details, such as animal ID, day, session stage. Select the data set to be explored by clicking the corresponding window of the footprint, then click the “cell exploration” button to open a separate window that is subdivided into five main components (**Fig. 2**), including

1. Ca^2+^ imaging video: toggle between original *vs*. CNMF processed videos, can be zoomed in to see more details about the cellular signal.
2. Behavioral video tracking, showing behaviors of the freely-moving animal during Ca^2+^ imaging recording,
3. Cell list, including individual cell number identified by CNMF,
4. Ca^2+^ trace window,
5. Trace toolbox

**Figure 1.**
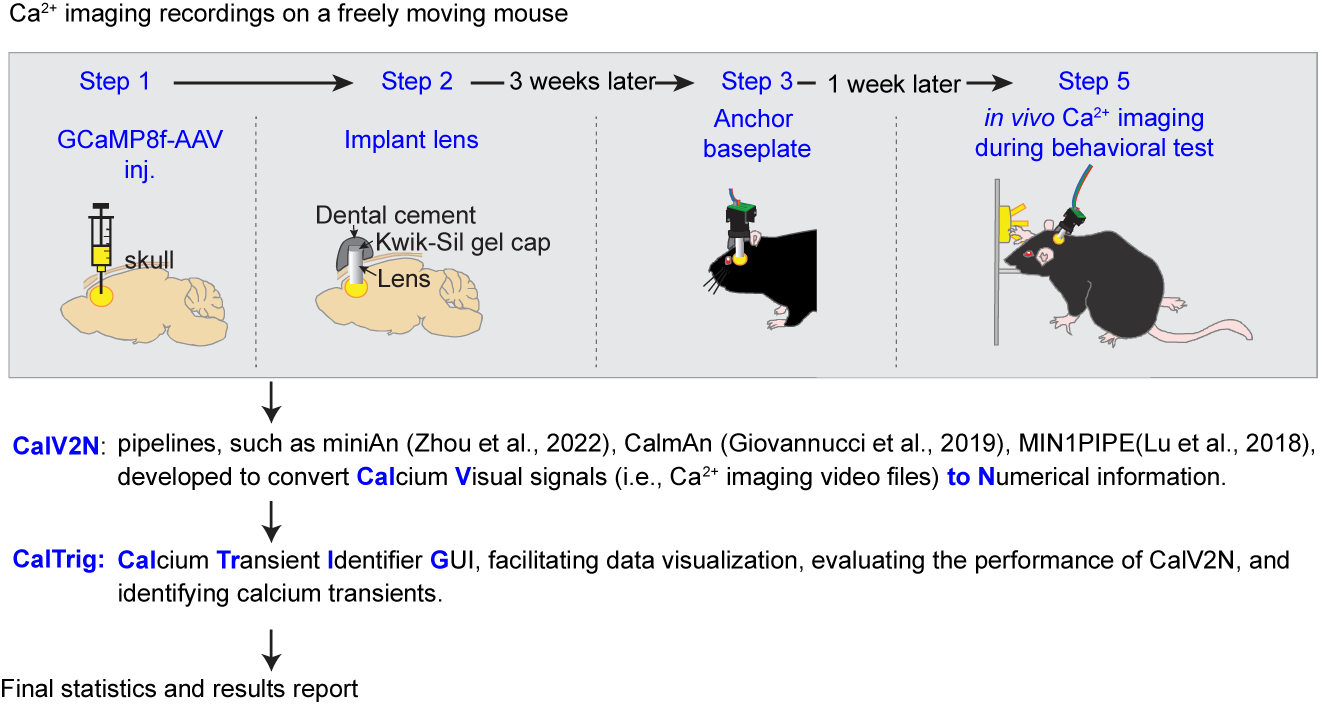
Flowchart showing the procedures of an *in vivo* Ca^2+^ study.

**Figure 2.**
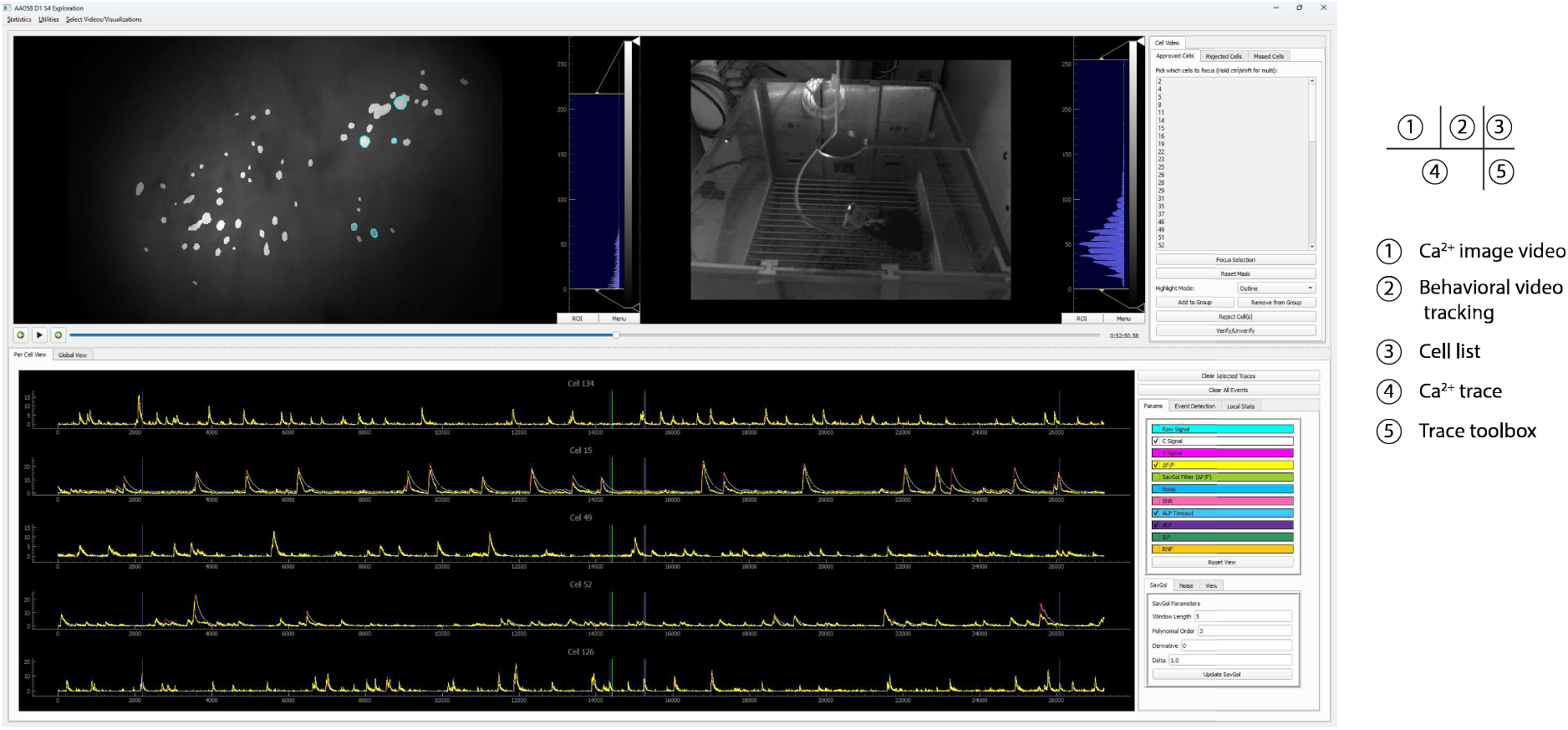
Five windows in CalTrig interface.

The following tasks can be done by the crosstalk between windows:

- <u>Footprint of cells of interest</u>: The footprint of the detected cells can be visualized by (1) clicking on the projection area of a specific cell or selecting an image region containing multiple cells in the Ca^2+^ imaging video, or (2) choosing the cell number from the cell list. This action superimposes its corresponding footprint onto the Ca^2+^ imaging video. The footprint can be displayed as a solid patch, a contour or by dimming the non-cell intensity.
- <u>Ca^2+^ traces of cells of interest:</u> Display the Ca^2+^ transient traces of the cells of interest, which can be selected by clicking on the footprint area in Ca^2+^ imaging video, or selection of the cell number in the cell list.
- <u>Variable readouts of Ca^2+^ traces:</u> Ca^2+^ traces can be visualized with different temporal readouts as listed in Trace toolbox, including the variables directly imported from CNMF (i.e., C signal and S signal), the variables calculated by CalTrig (i.e., ΔF/F, Raw signal, SavGol Filter of ΔF/F, Noise, SNR).
- <u>Time stamp of behavioral events or external stimuli on Ca^2+^ traces:</u> Ca^2+^ traces can be segmented by the behavioral readout listed in the Trace toolbox, such as ALP, ILP, RNF.
- <u>Standardized window size of the Ca^2+^ trace window:</u> The window size for X-axis in the unit of frame, and the scale range for Y-axis, can be defined in this box. When clicking “Reset view” button, the magnification of the signal is adjusted according to the Y axis range, and the window is set up with the pre-defined size by anchoring the first visible frame in the Ca^2+^ trace window. This is considerably helpful when manually identifying or verifying Ca^2+^ transients.
- <u>Cell verification</u>: Cells identified by CNMF can be individually reviewed and categorized into “Approved cells”, “Rejected Cells” and Missed Cells” as detailed below by crosstalk between multiple windows.
- <u>Quality evaluation of CNFM analysis:</u>
- <u>Ca^2+^ transient identification:</u> this can be done by parameter-based auto detection, manual detection, machine learning-based detection, or combination of multiple strategies we have developed in CalTrig.

#### 2.3.3. Cell verification and CalV2N evaluation

##### Cell verification

- <u>Verified cell:</u> CalTrig allows user to inspect individual cells detected by CalV2N by interactive exploration of the Ca^2+^ imaging window and Ca^2+^ trace window. A cell becomes verified after a visual confirmation of its footprint, Ca^2+^ Image, and Ca^2+^ trace, ensuring identifiable Ca^2+^ transients are present (**Fig. S3**). Once verified, the cell can be analyzed using tools developed for Ca²⁺ transient identification, as outlined in **Figs. 3, 4**.
- <u>Rejected Cell</u>: Cells can be assigned to the rejection column for different reasons, including no identifiable Ca^2+^ transients, suspicious footprint, statistically identified as an outlier of the detected Ca^2+^ transients, etc. CalTrig provides the option to provide a written justification for each cell rejection, which can be used to provide feedback and potentially rerun the CNMF pipeline.
- <u>Missing Cell</u>: Automatic time-series based algorithms may not be sensitive enough to detect cells with low to medium SNR, or those constitutively active or inactive with minimal dynamic changes. While reviewing the Ca^2+^ imaging video, one may notice a potentially missed cell by CalV2N. CalTrig allows user to manually draw the contour of the suspected cell in the Ca^2+^ imaging window, creating its footprint (**Fig. S4**). The corresponding temporal trace of signal intensity for the selected area is then calculated by averaging pixel intensities and displayed in the Ca^2+^ trace window. If the Ca^2+^ traces meet the criteria for accepted cells, the manually identified cell can be added to the list of missing cells.

**Figure 3.**
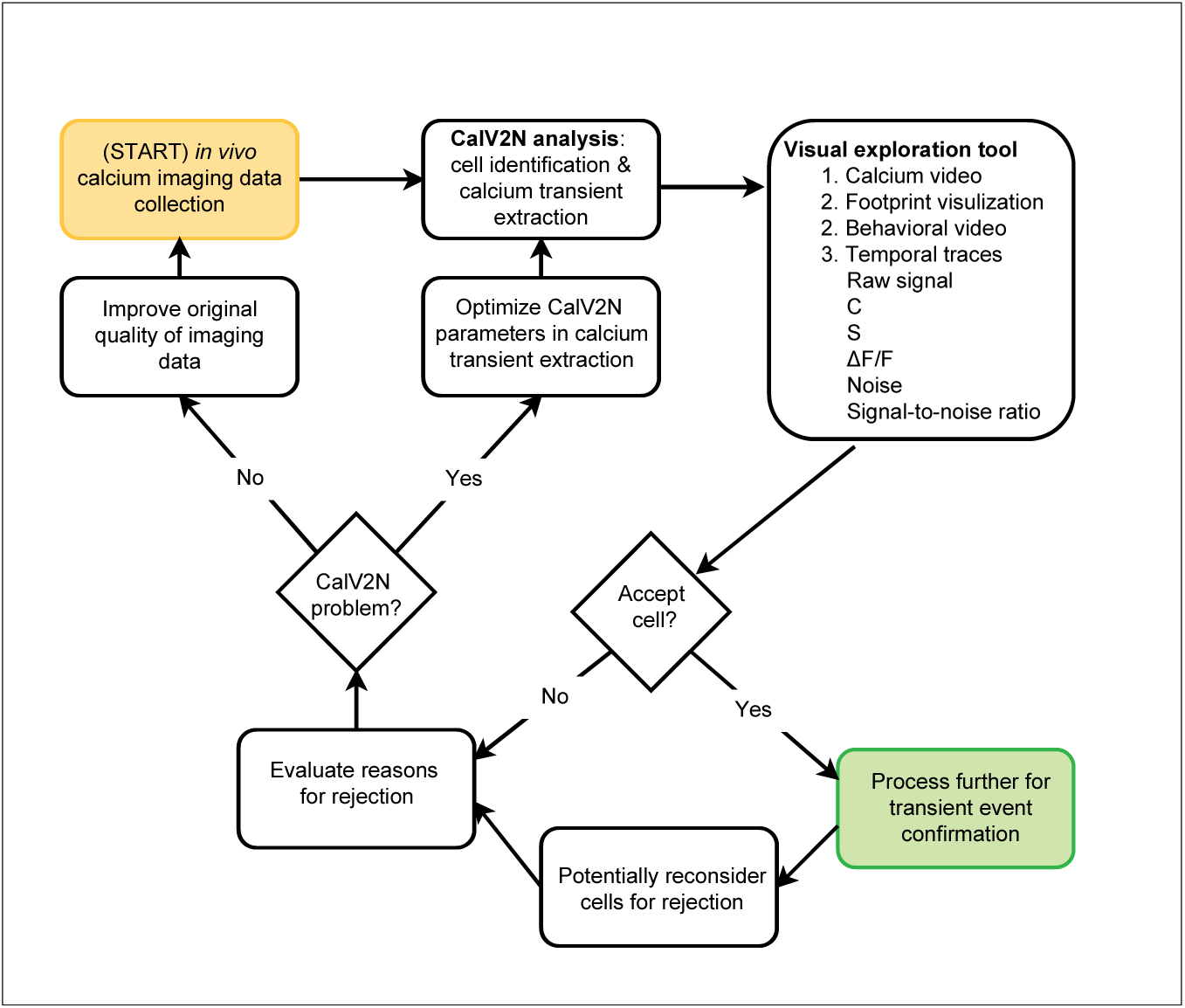
Use CalTrig to validate and filter cells identified by CalV2N.

**Figure 4.**
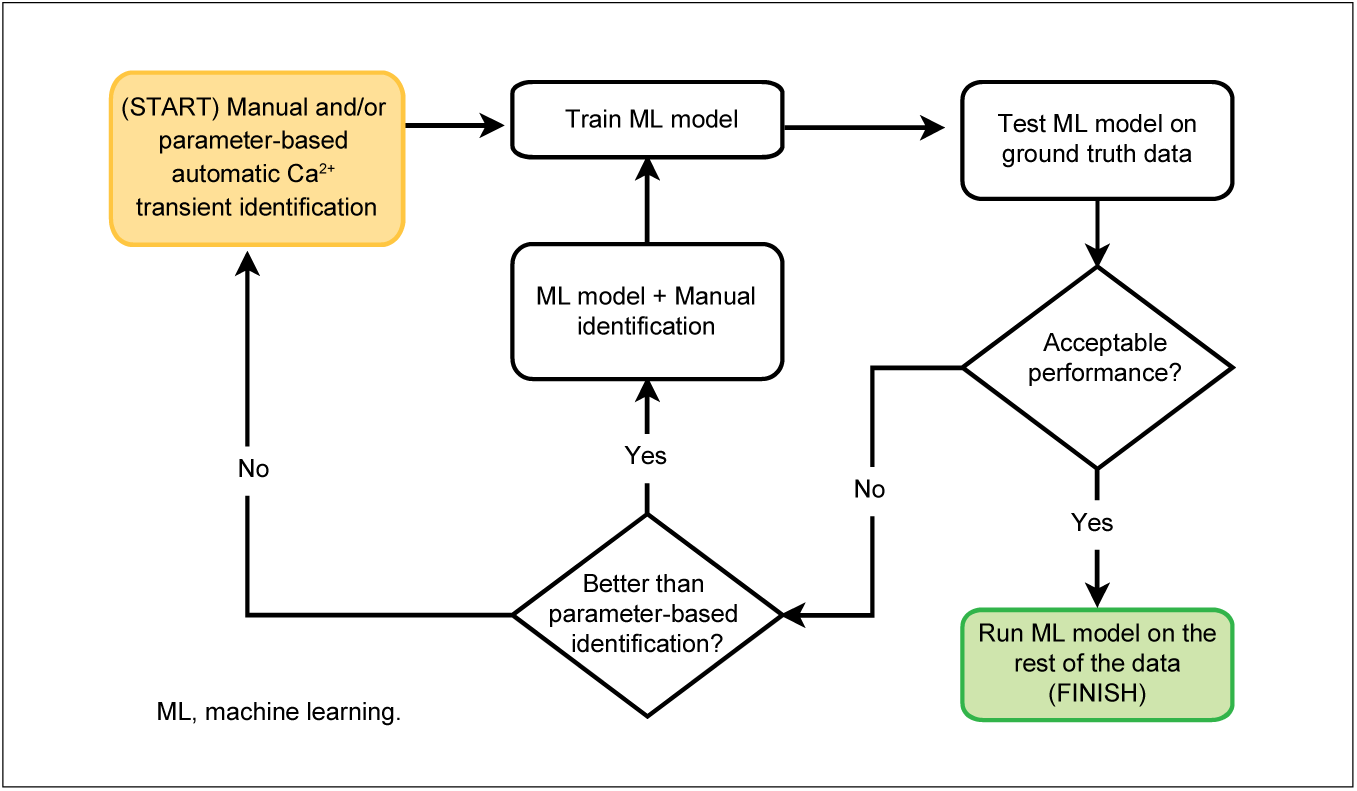
Use CalTrig to identify Ca^2+^ transients.

##### Evaluation of CalV2N performance

Although many CalV2N tools have been developed, there is no evaluation system available to assess the quality of cell identification and Ca^2+^ transient extraction. One of our initial motivations for developing CalTrig was to address this gap by creating a platform to evaluate CalV2N outputs (**Fig. 3**). Cells identified by CalV2N can be accepted or rejected, and missing cells can be manually added as described above. Further refinement of cell quality evaluation is possible after Ca^2+^ transient detection, using metrics such as rise time, peak amplitude, and inter-transient interval to review transient kinetics. If the rate of missing cells is too high (e.g., 10% or higher) and/or the rate of acceptable cells is too low (e.g., 90% or lower), further optimization of CalV2N parameters should be considered. For example, MiniAn pipeline can be optimized by adjusting the following parameters^11^.

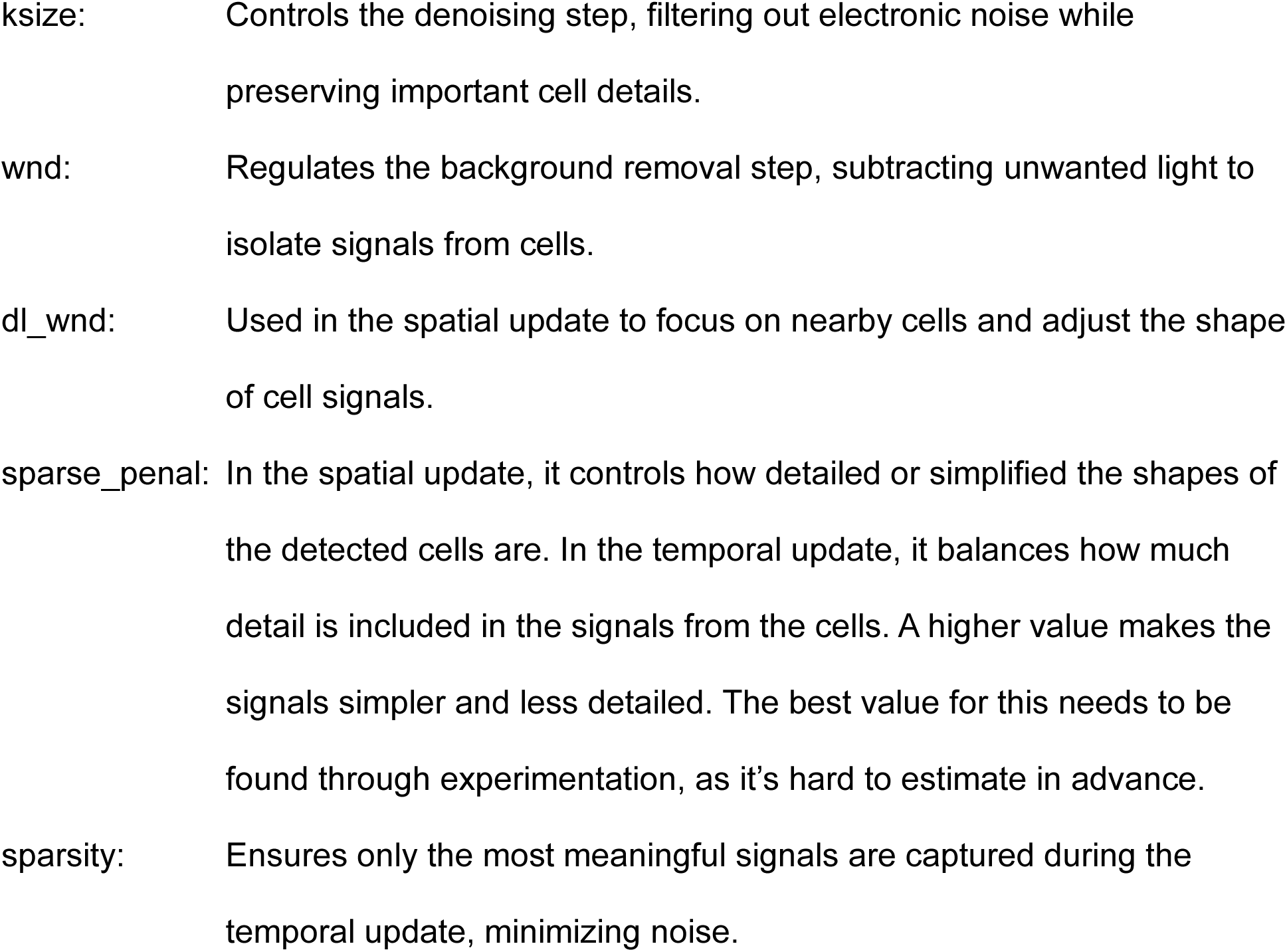

Low SNR may also indicate poor raw data quality, which can be addressed by optimizing AAV preparation (e.g., selecting appropriate subtypes, adjusting titer and volume), improving surgical techniques (slower needle insertion / withdrawal, slower AAV delivery, minimizing bleeding), and enhancing data collection methods (adjusting focal levels, securing MiniScope anchoring, preventing cable twists *via* sensitive commutator, and ensuring reliable hardware connections).

#### 2.3.4. Ca^2+^ transient identification

Transient confirmation can be done in three ways, including manually selecting the start and end points of the transients, auto filtering with specified kinetic parameters or specifying frame number ranges, or running a trained machine learning model.

##### 2.3.4.1. Manual Identification

<u>Method</u>: Ca^2+^ transients can be directly identified by manually selecting the start and end points of the rising section of Ca^2+^ transients (**Fig. S5**). Efforts have been made to improve the efficiency of manual identification. For example, the window size of the Ca^2+^ can be pre-set at a desired size (e.g., 1000 frames per window width in the x-axis, −1 to 20 as the signal range in the y-axis), which will allow the visual impression of Ca^2+^ at different time stage, different animals are comparable, assisting a consistent decision making on Ca^2+^ manual identification. We also set up the shortcut keys to switch the screen window in the Ca^2+^ trace panel. Specifically, clicking A and F allows to jump to the first or the last window, respectively, and clicking S and D allows to jump to the previous or the next screen window, respectively. To account for inaccuracies resulting from human interaction, the selected points would automatically position themselves within a 20-frame window to the local maxima or minima, dependent on their order.

<u>Applications</u> for manual identification: **First**, directly label the Ca^2+^ transient at the beginning stage of the project when no reference in setting up the parameters for autodetection or no data set are available to training the machine learning model. It takes ∼5 min to go through a 15-min Ca^2+^ traces. **Second**, correct, add, or remove the transient spikes after running the parameter-based auto detection to establish the ground truth for training the machine learning model. This would primarily apply to problematic spikes around the specified parameter threshold or those whose rising part is too slow to be confidently identified as a transient event. It takes ∼1 min to go through a 15 min Ca^2+^ traces. **Third**, to further improve the Ca^2+^ transient identification after processed by an established machine learning model. Since our machine learning model, specifically the GRU model, used in identifying the Ca^2+^ transient in neurons detected during the same recording window from the same animal are highly reliable (more details in **Results**), we expect this correction process should be done on 2-4 cells per min. **Fourth**, when we need to extend the application of a well-trained machine learning model to a different brain region, a different cell type, *etc*., the machine learning model may provide a compromised predictability in detecting Ca^2+^ transients. Adding a few traces with the manually identified Ca^2+^ transient will set up a new data set as ground truth for establishing an updated machine learning model to be used in a new task.

##### 2.3.4.2. Parameter-based automatic identification

###### Three Parameters

“Peak Threshold (ΔF/F)”: First, let’s define ΔF/F, which represents the relative change in fluorescence intensity, where:

- **F₀** is the background fluorescence intensity, which is calculated using a moving percentile provided by Caiman^4^.
- **ΔF** is the difference between the observed fluorescence intensity (**F**) at a given time and the background fluorescence intensity (**F₀**).

The formula is:

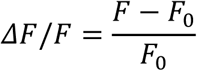

The peak ΔF/F value within a specified time window is considered as the potential peak of the Ca^2+^ transient. The “Peak Threshold (ΔF/F)” is the minimum value of peak ΔF/F. A Ca^2+^ transient is accepted only if its peak ΔF/F surpasses this threshold.

“Interval Threshold”: This specifies the minimum Inter-Transient Interval (ITI), measured as the frame distance between the initial fames of two adjacent Ca^2+^ transients. If the distance between two transients is shorter than the “Interval Threshold”, they are considered part of the same transients, with the lower peak, whether preceding or following, merged into the higher peak.

“SNR Threshold”: This sets the minimal Signal-to-Noise ratio (SNR). Using the Savitzky–Golay filter ^12,13^ to smooth ΔF/F signals, noise is calculated as the difference between the original and filtered ΔF/F, further smoothed by a rolling window strategy. Then SNR is computed by dividing the smoothed ΔF/F by the estimated noise. See more details in **Supplementary Information** and **Fig. S6**.

At the initial experimental stage, parameter settings are determined using manual identification on a limited number of cells or demo data. As more cells are manually identified, these parameters can be refined to enhance the effectiveness of parameter-based auto-detection.

<u>Method</u>: Through manual identification, the C signal from CNMF was found to reliably predict Ca^2+^ transients in most cases but prone to false positives. To enhance detection accuracy, the parameter-based auto detection algorithm starts by including all C peaks in a candidate pool, then filtering down to a valid subset based on three pre-defined parameters: “Peak Threshold (ΔF/F)”, “Interval Threshold”, and “SNR Threshold” (**Fig. S7**). This is achieved through the following steps.

- Initial peak detection through local maxima calculation of the C signal.
- Refine peak selection from left to right by looping the following steps.
- Remove erroneous peaks if the S signal is zero.
- Allocate overlapping, continuous S signal to the current peak.
- Check if the distance to the next peak is shorter than the pre-defined “<u>Interval Threshold</u>”.
- If so, merge the current and subsequent peaks, selecting the taller peak.
- If not, use the S signal to determine the start and end of the transient.
- Accept the transient selection if in the defined boundaries, the values surpass both the pre-defined “Peak Threshold (ΔF/F)” and “SNR Threshold”.

<u>Application</u>: Parameter based Ca^2+^ transient detection is faster than manual detection, but less predictive than using a well-trained machine learning model. It is particularly feasible after manual identification of a few cells, enabling parameter tuning. Parameter-based detection can then serve as a pre-detection step to accelerate further manual identification, eventually creating a ground-truth dataset for training machine learning models. This method also helps refine or update machine learning models for different experimental conditions (**Figs. 6, 8, 9**).

##### 2.3.4.3. Detection using Machine Learning Model

###### Feature selection for machine learning model

In selecting the appropriate architecture for the machine learning model, we aimed to incorporate key signals gained from the “ground truth” Ca^2+^ transient verification by manual detection process. We identified two signals, i.e., C and ΔF/F, as sufficient for decision-making. The determination of whether a given time-step contains a Ca^2+^ transient can be inferred from a 100-frame window centered by the frame of interest. A corresponding data set, denoted E, was created by marking the initial and peaking frames within the context of a dynamic rise in Ca^2+^ transients.

###### Selection of machine learning models

Ca^2+^ transient data are inherently time-series data with a long sequence of many neurons firing at different times, where the signal at any given frame is highly relevant to past and upcoming activity. Thus, Recurrent Neural Networks (RNNs) ^14^ and Transformers^15,16^ are selected as the candidate machine learning models for Ca^2+^ transient detection due to their ability or potential to handle the temporal dynamics of Ca^2+^ signals in neurons (**Fig. 5**).

**Figure 5.**
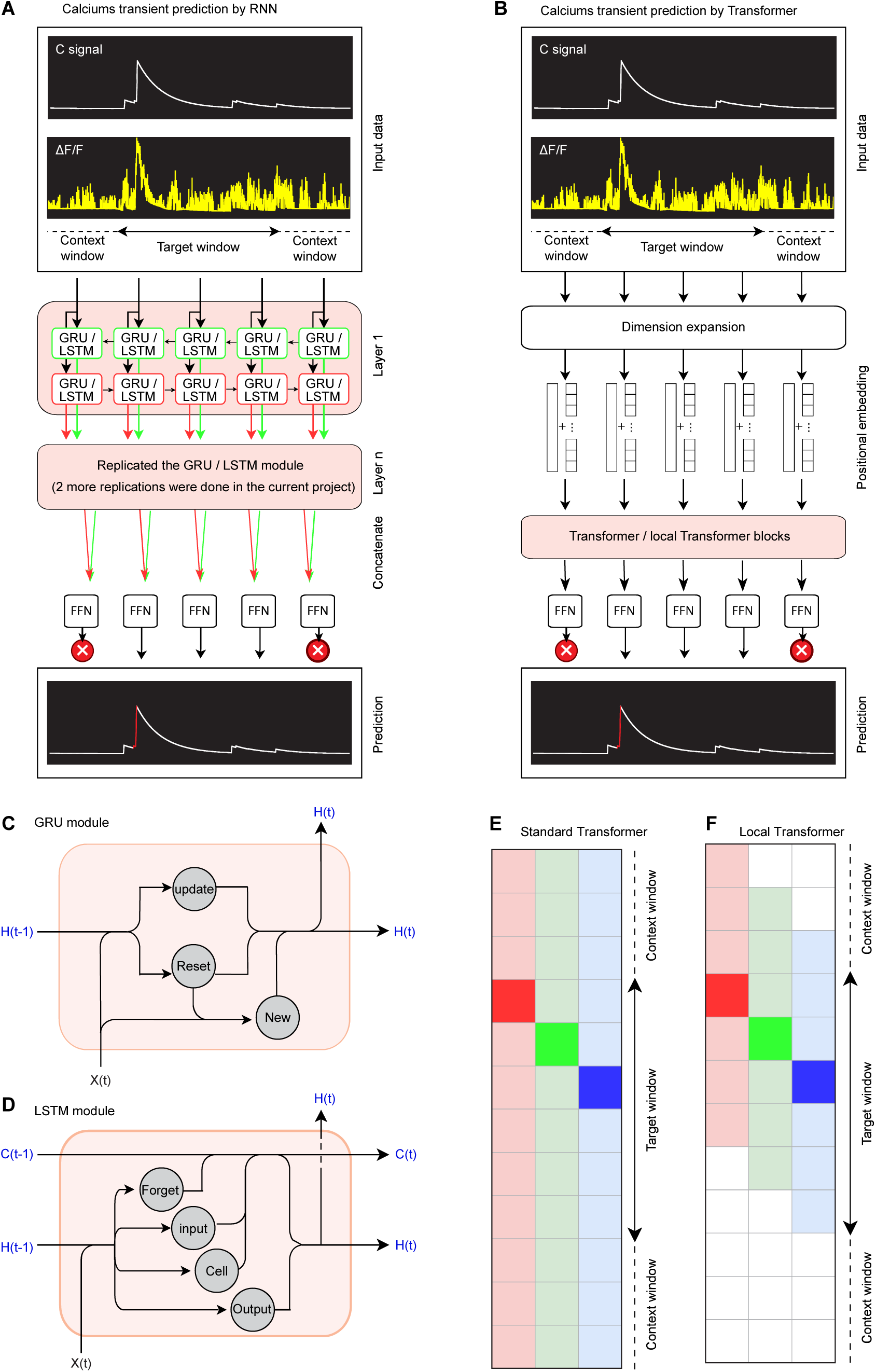
Flowchart demonstrating Ca^2+^transient validation via machine learning models. **A**, Applications of RNN in predicting Ca^2+^ transients. **B**, Applications of Transformer in predicting Ca^2+^ transients. **C**, **D**, Diagrams showing the internal operations of two variants of RNN, i.e., GRU module (**C**) and LST module (**D**). The GRU uses two gates: the *update gate*, which controls the amount of information passed to the next step, and the *reset gate*, which determines how much of the previous information to forget. These gates modify the hidden state H(t) at the current time step based on the input X(t) and the previous hidden state H(t−1), leading to a new hidden state H(t). The LSTM module architecture is depicted, showcasing its cell structure. It uses three gates: *forget*, *input*, and *output* gates to control the flow of information. The input X(t), the previous cell state C(t−1), and the previous hidden state H(t−1) are processed through these gates to update the cell state C(t) and produce the new hidden state H(t). **E**, **F**, Diagram showing the Transform (**E**) and local Transformer (**F**) mechanisms. Each column corresponds to the same set of data. The non-translucent color in a column highlights the current input being evaluated, while translucent colors indicate the values being compared to it. No color signifies that certain values are excluded from evaluation for the current input. FNN, Feedforward neural network.

The core of its architecture lies in the RNN cells and its gating mechanisms, which processes inputs sequentially, creating a representation of the data in the context of the preceding timestamps. In the case of Ca^2+^ transient data, two features (i.e., C, ΔF/F) used for training will be incorporated into the hidden state (*H*) and passed to the subsequent cell (*C*). We tested two RNN variants: Long Short-Term Memory (LSTM) ^17,18^ and Gated Recurrent Unit (GRU)^19,20^. Given the importance of both forward and backward context in detecting Ca^2+^ transients, we employed a bidirectional RNN. The outputs from both directions are concatenated and passed through a feed-forward layer to generate the final prediction.

One significant limitation of the RNN is due to the fixed size of its hidden state and the sequential nature of the architecture, causing early-time step bias. To address this, we trained a Transformer, which uses self-attention to compare the relationship between all input values simultaneously, rather than processing sequentially^21,22^. This approach allows the model to attend to both low and high-intensity values that occur at different temporal distances from the Ca^2+^ transient, improving prediction accuracy. Recent advances in Natural Language Processing have introduced the **Local Transformer**, which limits attention to a subset of nearby inputs ^23,24^. This model could help reduce errors by focusing on local context and ignoring distant irrelevant signals. However, the self-attention mechanism in the Transformer obscures positional information, hence we need positional embeddings. The trainable embeddings were used in our Transformer. Additionally, we observed that the small dimensionality of the original input data negatively affected training, likely due to the model’s inability to effectively encode positional information in low-dimensional data. To address this, we introduced a preliminary dimension expansion layer, which projects the input into a higher-dimensional space before passing it through the encoding layers.

Both RNN and Transformer architectures were implemented in CalTrig using PyTorch ^25,26^. For the Transformer, we used the Transformer Encoder module, and for the Local Transformer, we adapted a modified version of the model ^27^. Parameters for machine learning model training are listed in **Table 2**.

**Table 2.** Parameters for machine learning model training.

| Parameters | GRU & LSTM | Transformer | local Transformer |
| --- | --- | --- | --- |
| Hidden size | 30 | 42 | 32 |
| Number of layers | 3 | 3 | 3 |
| Loss function | BCEWithLogitsLoss | BCEWithLogitsLoss | BCEWithLogitsLoss |
| Optimizer | Adam | Adam | Adam |
| Initial learning rate | 0.001 | 0.001 | 0.001 |
| Section Length | 200 | 200 | 200 |
| Slack | 50 | 50 | 50 |
| number of Heads |  | 2 | 1 |
| Local Attention Window Size |  |  | 10 |
| Look forward/backward Size |  |  | 5 Local Attention Windows |

###### Data Pre-processing

The input data consisted of two signals (i.e., C, ΔF/F) representing 15-minute time intervals of 27,000 frames per cell, captured at a sampling rate of 30 Hz. Each segment was normalized relative to the highest value within the cell, per data type. We tested two data generation and loading approaches:

Discrete Sample Chunking: We defined three variables—sequence length, slack, and rolling parameter. The sequence length specifies the number of frames the model is trained on to make predictions, typically set to around 200 frames in our testing. The slack variable determines the extra context length on either side of a given sequence, providing necessary context for making predictions without being used for the predictions themselves. This slack is usually set between 50 and 100 frames. For sequences at the edges of the 27,000-frame segment, zero-padding is applied to match the slack length and maintain consistency. Lastly, the rolling parameter defines the windowing approach for generating data, allowing overlapping sequences to be extracted for model training.

Ca^2+^ transient events account for only 2-3% of the overall data. To avoid biasing the model towards detecting non-Ca^2+^ transients, we initially applied class weighting, which resulted in a poor precision score. We determined that it was due to the underemphasizing of noisy data within training, whose signal characteristics were more similar to transient activity rather than an empty signal. This resulted in a model that considered any activity including noise to be a transient event. We opted instead to implement stratification where we ensured that only samples containing ground-truth transient events or positive values from the C array were included. The C array was used as a reference to give the model insight into problematic spikes identified by the CNMF process but deemed invalid by the verifier. During classification, we average all outputs for a single time-step to address overlapping predictions.

###### Key metrics for validation

We used Precision, Recall, F1 and macro F1 as key metrics to evaluate the performance of machine learning model in predicting Ca^2+^ transients *vs*. non-Ca2+ transients (**Fig. S8**). There are four types of predictions as shown in **Tables 3, 4**: true positive (TP) predicting the positive as positive, true negative (TN) predicting the negative as negative, false positive (FP) predicting the negative as positive, and false (FN) negative predicting the positive as negative.

**Table 3.**
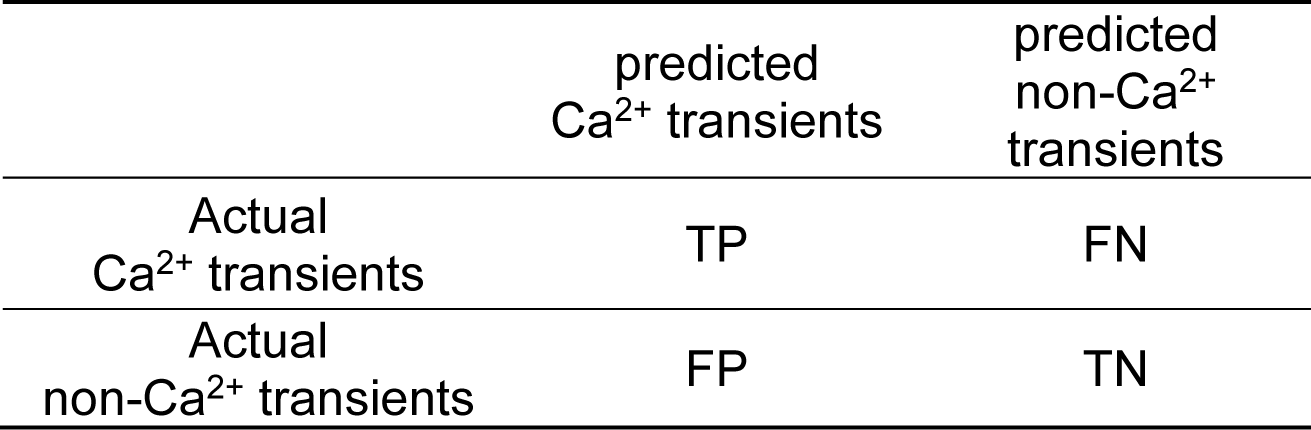
Event to be predicted: Ca^2+^ transients.

**Table 4.** Event to be predicted: non-Ca^2+^ transients.

| | predicted<br>$\text{Ca}^{2+}$<br>transients | predicted<br>non-$\text{Ca}^{2+}$<br>transients |
| --- | --- | --- |
| Actual<br>$\text{Ca}^{2+}$ transients | TN | FP |
| Actual<br>non-$\text{Ca}^{2+}$ transients | FN | TP |

Precision measures how many of the positive predictions (TP + FP) are actually positive (TP). (It focuses on the quality of the positive predictions.)

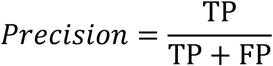

Recall measures how many of the actual positive instances (TP + FN) are correctly predicted as positive (TP). (It focuses on the ability to find all positive cases.

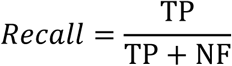

F1 score is the harmonic mean of Precision and Recall, balancing both metrics.

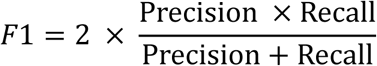

Macro F1 is the average of the F1 scores calculated for each class, i.e., Ca^2+^ transients and non-Ca^2+^ transients.

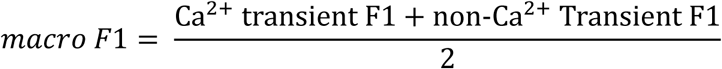

For evaluation, cells were randomly separated into training (80%), validation (10%), and testing (10%) sets (**Fig. 6A**). Manual identification, parameter-based identification and machine learning model-based identification can be integrated in detecting Ca^2+^ (**Fig. 4**).

**Figure 6.**
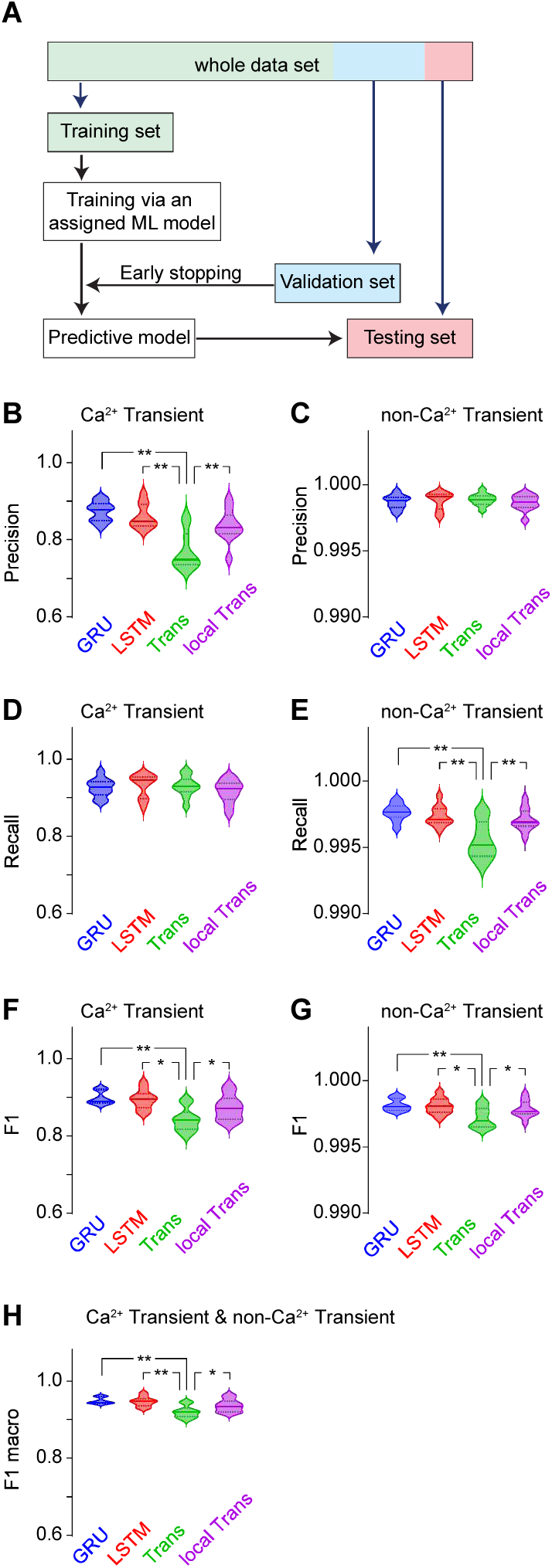
GRU module has the best predictability of Ca^2+^ transients. **A**, Shared strategy for assigning data to training, validation, and testing across four machine learning models. **B**, **C**, Precision varied significantly in Ca^2+^ transient prediction (**B**, F_3,36_=14.2, p<0.01), but remained similar in no Ca^2+^ transient prediction (**C**, F_3,36_=0.4, p=0.78) in four machine learning models. **D**, **E**, Recall remained similar in Ca^2+^ transient prediction (**D**, F_3,36_=0.6, p=0.63) but varied in Ca^2+^ transient prediction (**E**, F_3,36_=9.4, p<0.01) in four machine learning models. **F**, **G**, Significant differences in F1 scores of Ca^2+^ transient prediction (**F**, F_3,36_=9.3, p<0.01) and no Ca^2+^ transient prediction (**G**, F_3,36_=4.8, p<0.01) in four machine learning models. **H**, Significant differences in macro F1 scores (F_3,36_=9.3, p<0.01) in four machine learning models. Each machine learning model was trained using data from 226 cells, with 28 cells allocated to the validation set and 28 cells to the testing set. Data were analyzed by one-way ANOVA, followed by Bonferroni *post hoc* test. *, p<0.05; **, p<0.01.

#### 2.3.5. Data export

All the detected Ca^2+^ transient information can be extracted. A few examples are listed below. This will directly aid in statistical analyses and figure preparation for sharing and publishing.

##### Animal-wide data export

A data table is generated, with each row representing one of the CalS2N-detected cells and each column providing specific readout as a general description of that cell (**Fig. S9A**). From left to right, the columns include: Cell ID, Cell Size (number of pixels), Footprint Location (x, y), Total Ca^2+^ Transient Count, Frequency (Hz), Average Amplitude (ΔF/F), Average Rising (# of frames), Average Rising Time (seconds), Average Interval (seconds), Standard Deviation (denoted Std, ΔF/F), Mean Absolute Deviation (denoted MAD, ΔF/F), Average Peak Amplitude (ΔF/F), and Category (Verified, Rejected, Missing). The data can be easily copied to the clipboard for further processing in other applications like Excel (Microsoft 365) or Prism (GraphPad). Figures, such as box plots showing the 25%, 50%, and 75% values for metrics like Average Peak Amplitude or ITI (**Fig. S9B, C**), can be directly generated and saved as editable SVG files for publication purposes.

##### Cell-wide data export

A data table is generated for a selected cell, with the top row displaying column titles and the subsequent rows representing individual Ca^2+^ transients detected by CalTrig, one transient per row (**Fig. S10A**). The columns include: Rising Start (frames), Rising Stop (frames), Total Rising Frames, Rising Start (seconds), Rising Stop (seconds), Total Rising Time (seconds), Interval with Previous Transient (frames), Peak Amplitude (ΔF/F), and Total Amplitude (ΔF/F, calculated as the area under the rising slope). The column title can be selected to sort the data based on the information in the selected column (**Fig. S10B**). This table can be copied to the clipboard for further processing in applications like Excel (Microsoft 365) or Prism (GraphPad). Figures such as Amplitude Distribution or ITI Frequency Histograms (**Fig. S10 C, D**) can be created directly from this data and saved as editable SVG files for publication.

##### Maximum projection image

An editable image file, including the footprints of detected or verified cells for an animal, can be created for data sharing and publication purposes.

### 2.4. Statistical Analysis

Data were collected from 10 mice *in vivo,* shown as mean ± SEM in curve graphs (**Figs. 8B, C, F, G, J, K, N; 9B, C, F, G, J, K, N**) or the quartile in violin plots (**Figs. 6; 7; 8D, E, H, I, L, M, O; 8D, E, H, I, L, M, O**). Using GraphPad Prism 10, statistical significance was assessed by one way ANOVA or two-way ANOVA, followed by Bonferroni post-hoc tests. Statistical significance was considered to be achieved if p < 0.05.

**Figure 7.**
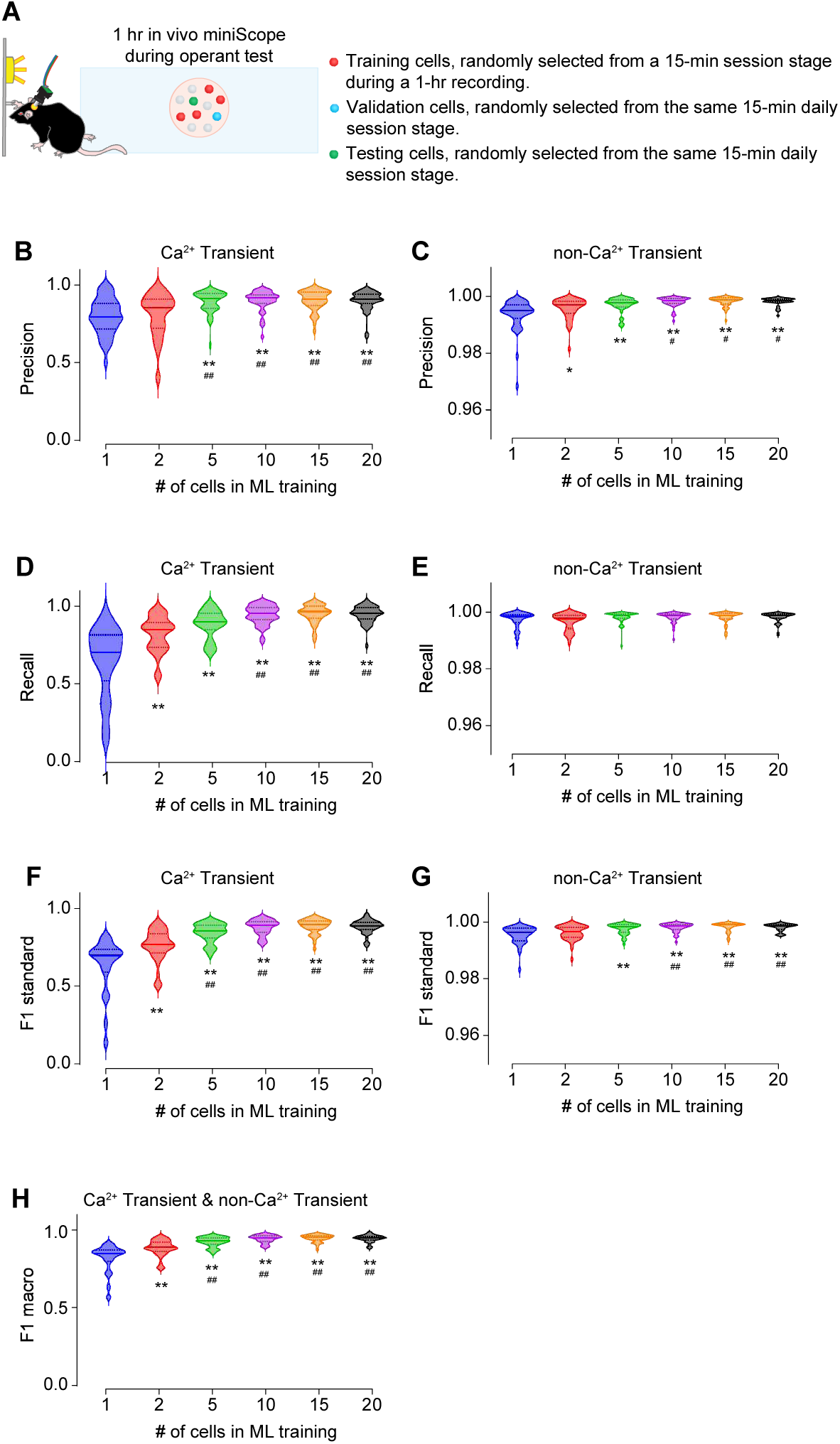
Prediction of Ca^2+^ transients or no Ca^2+^ transients across different numbers of cells in machine learning training. **A**, Strategy of randomly picking up cells for training, validation and testing within the same session stage on the same day. **B**, **C**, Increasing the number of cells in machine learning training model significantly improved the precision in predicting Ca^2+^ transients (**B**, F_5,234_=11.0, p<0.01) and no Ca^2+^ transients (**C**, F_5,234_=12.8, p<0.01). **D**, **E**, Increasing the number of cells in machine learning training model significantly improved the recall in predicting Ca^2+^ transients (**D**, F_5,234_=40.9, p<0.01) and no Ca^2+^ transients (**E**, F_5,234_=2.8, p=0.02). **F**, **G**, Increasing the number of cells in machine learning training model significantly improved the F1 scores in predicting Ca^2+^ transients (**F**, F_5,234_=46.7, p<0.01) and no Ca^2+^ transients (**G**, F_5,234_=12.0, p<0.01). **H**, Increasing the number of cells in machine learning training model significantly improved the macro F1 scores in predicting Ca^2+^ transients and no Ca^2+^ transients (**H**, F_5,234_=46.6, p<0.01) Data were analyzed by one-way ANOVA, followed by Bonferroni *post hoc* test. *, p<0.05; **, p<0.01, compared to the machine learning model trained by 1 cell. ^#^, p<0.05; ^##^, 0.01, compared to the machine learning model trained by 2 cells. 40 testing cells in each group.

**Figure 8.**
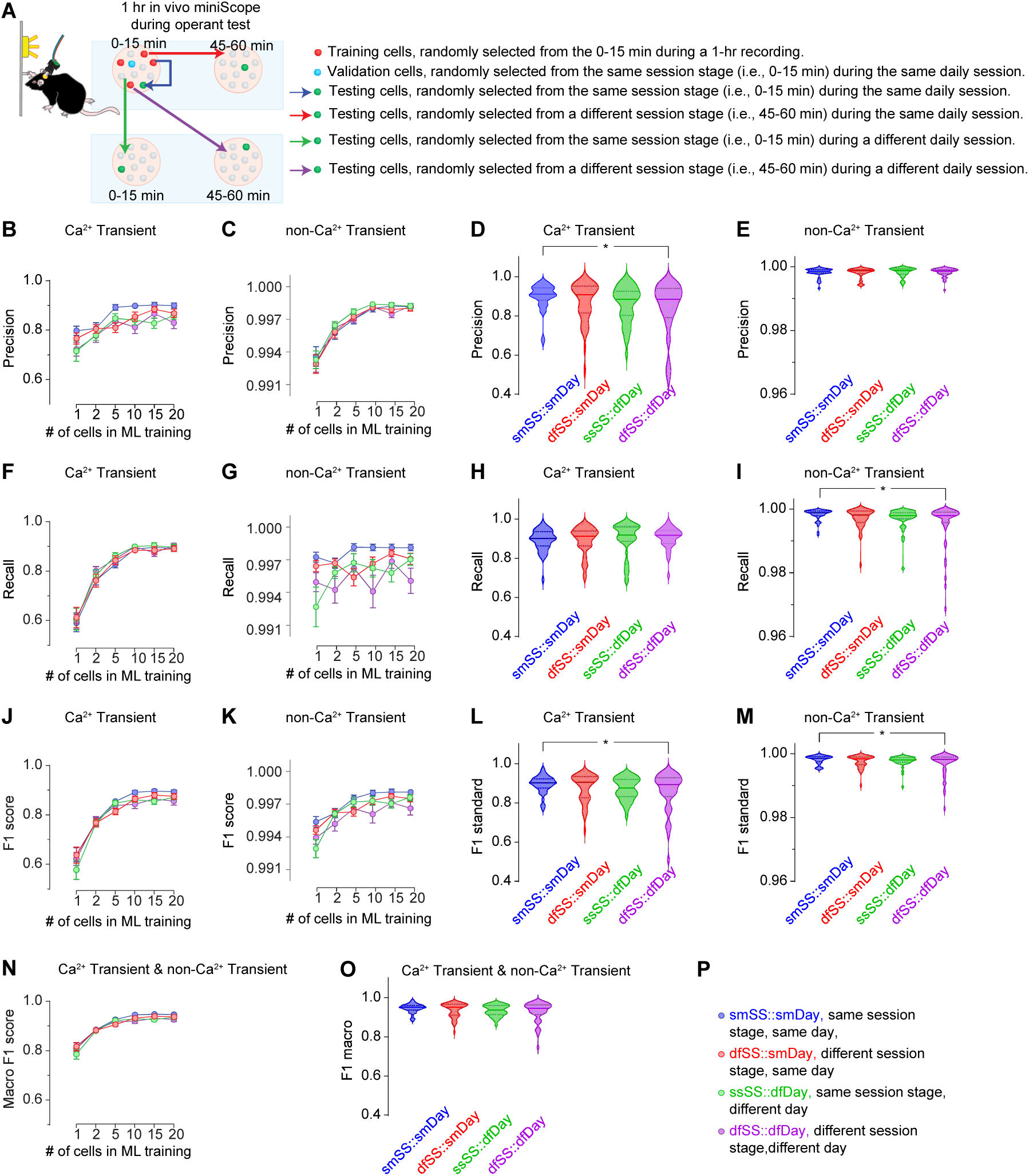
Prediction of Ca^2+^ transients or no Ca^2+^ transients within the same mice. **A**, Four strategies for sourcing cells within the M2 area from a mouse for machine learning model training, validation, and testing. **B-E**, The training cell number, and the testing cell source, but not their interactions, significantly affected the Precision in predicting Ca^2+^ transients (**B**, training cell # F_5,936_=21.4, p<0.01; testing cell source F_3,936_=10.0, p<0.01; training cell number × testing cell source interaction F_15,936_=0.8, p=0.64). The training cell number, but not the testing cell source or their interactions, affected the Precision in predicting no Ca^2+^ transients (**C**, training cell number F_5,936_=71.7, p<0.01; testing cell source F_3,936_=1.1, p=0.36; training cell number × testing cell source interaction F_15,936_=0.3, p=0.99). When 20 cells were included in the machine learning training model, the Precision in predicting Ca^2+^ transients (**D**, F_3,156_=2.7, p=0.046), but not in predicting no Ca^2+^ transients (**E**, F_3,156_=0.2, p=0.93), in testing cells from different session stage on different day reduced. **F-I**, The training cell number, but not the testing cell source or their interactions, significantly affected the Recall in predicting Ca^2+^ transients (**F**, training cell number F_5,936_=126.0, p<0.01; testing cell source F_3,936_=0.8, p=0.49; training cell number × testing cell source interaction F_15,936_=0.3, p=0.99). The number of cells, the testing cell source, but not their interactions, significantly affected the Recall in predicting no Ca^2+^ transients (**G**, training cell number F_5,936_=2.7, p=0.02; testing cell source F_3,936_=11.4, p<0.01; training cell number × testing cell source interaction F_15,936_=1.5, p=0.10). When 20 cells were included in the machine learning training model, the Recall in predicting no Ca^2+^ transients (**I**, F_3,156_=3.3, p=0.02), but not in predicting the Ca^2+^ transients (**H**, F_3,156_=0.4, p=0.78), in testing cells from different session stage on different day reduced. **J-M**, The training cell number and the testing cell source, but not their interactions, significantly affected the F1 scores in predicting Ca^2+^ transients (**J**, training cell number F_5,936_=130.0, p<0.01; testing cell source F_3,936_=3.0, p=0.03; training cell number × testing cell source interaction F_15,936_=0.7, p=0.74). Similarly, the number of cells and the testing cell source, but not their interactions, significantly affected the F1 scores in predicting no Ca^2+^ transients (**K**, training cell number F_5,936_=32.3, p<0.01; testing cell source F_3,936_=8.8, p<0.01; training cell number × testing cell source interaction F_15,936_=1.2, p=0.28). When 20 cells were included in the machine learning training model, the Recall in predicting Ca^2+^ transients (**L**, F_3,156_=2.7, p=0.048) and no Ca^2+^ transients (**M**, F_3,156_=2.8, p=0.040) in testing cells from different session stage on different day reduced. **N**, **O**, The training cell number, and the testing cell source, but not their interactions, significantly affected the macro F1 scores in predicting Ca^2+^ transients and no Ca^2+^ transients (**N**, training cell number F_5,936_=128.5, p<0.01; testing cell source F_3,936_=3.1, p=0.03; training cell number × testing cell source interaction F_15,936_=0.8, p=0.73). When 20 cells were included in the machine learning training model, the macro F1 scores in predicting Ca^2+^ transients and no Ca^2+^ transients (**O**, F_3,156_=2.8, p=0.041) in testing cells from different session stage on different day reduced. **P,** Legends showing the color-coded abbreviations for 4 experimental groups. Data were analyzed by two-way ANOVA (**B**, **C**, **F**, **G**, **J**, **K**, **N**) or one-way ANOVA (**D**, **E**, **H**, **I**, **L**, **M**, **O**), followed by Bonferroni *post hoc* test. *, p<0.05; **, p<0.01. 40 testing cells in each group.

**Figure 9.**
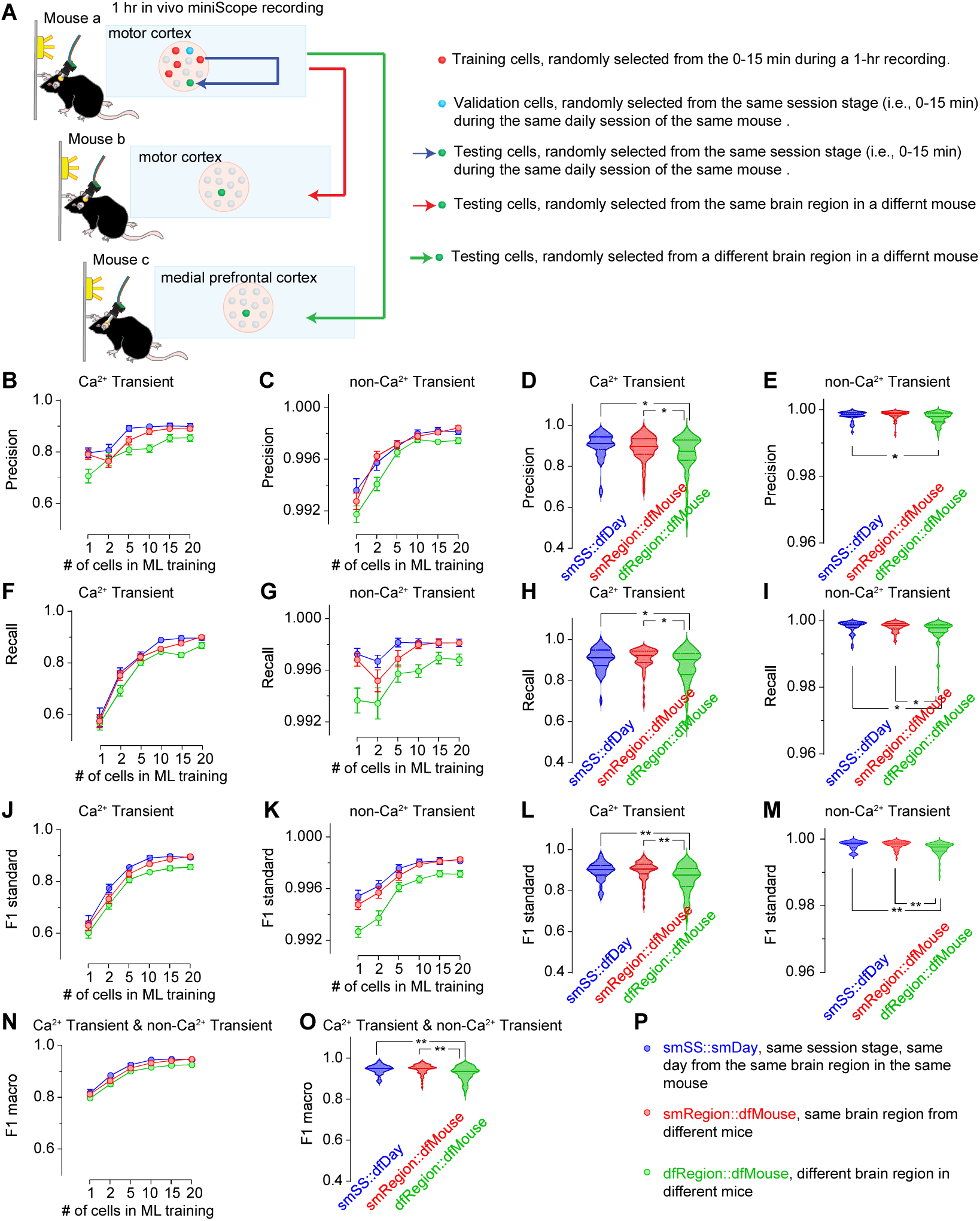
Prediction of Ca^2+^ transients or no Ca^2+^ transients in different mice. **A**, Three strategies for sourcing testing cells (smSS::smDay, between mice, different regions, see more details in panel **P**) **B-E**, The training cell number, and the testing cell source, but not their interactions, significantly affected the Precision in predicting Ca^2+^ transients (**B**, training cell number F_5,942_=24.4, p<0.01; testing cell source F_2,942_=19.2, p<0.01; training cell number × testing cell source interaction F_10,942_=1.1, p=0.37) and the no Ca^2+^ transients (**C**, training cell number F_5,942_=74.1, p<0.01; testing cell source F_2,942_=11.6, p<0.01; training cell number × testing cell source interaction F_10,942_=1.0, p=0.43). When 20 cells were included in the machine learning training model, the Precision in predicting Ca^2+^ transients (**D**, F_2,157_=4.7, p=0.01) and no Ca^2+^ transients (**E**, F_2,157_=6.2, p<0.001) in testing cells from different regions in different mice reduced. **F-I**, The training cell number, but not the testing cell source or their interactions, significantly affected the Recall in predicting Ca^2+^ transients (**F**, training cell number F_5,942_=145.6, p<0.01; testing cell source F_2,942_=3.7, p=0.02; training cell number × testing cell source interaction F_10,942_=0.7, p=0.68). The training cell number and the testing cell source, but not their interactions, significantly affected the Recall in predicting no Ca^2+^ transients (**G**, training cell number F_5,942_=8.7, p<0.01; testing cell source F_2,942_=22.9, p<0.01; training cell number × testing cell source interaction F_10,942_=0.9, p=0.58). When 20 cells were included in the machine learning training model, the Recall in predicting Ca^2+^ transients (**H**, F_2,157_=4.4, p=0.01) and no Ca^2+^ transients (**I**, F_2,157_=5.3, p<0.01) in testing cells from different regions in different mice reduced. **J-M**, The training cell number, and the testing cell source, but not their interactions, significantly affected the F1 scores in predicting Ca^2+^ transients (**J**, training cell number F_5,942_=165.7, p<0.01; testing cell source F_2,942_=17.6, p<0.01; training cell number × testing cell source interaction F_10,942_=0.3, p=0.98) and the no Ca^2+^ transients (**K**, training cell number F_5,942_=55.8, p<0.01; testing cell source F_2,942_=40.4, p<0.01; training cell number × testing cell source interaction F_10,942_=1.3, p=0.23). When 20 cells were included in the machine learning training model, the F1 scores in predicting Ca^2+^ transients (**L**, F_2,157_=8.9, p<0.01) and no Ca^2+^ transients (**M**, F_2,157_=9.3, p<0.01) in testing cells from different regions in different mice reduced. **N**,**O**, The training cell number, and the testing cell source, but not their interactions, significantly affected the macro F1 scores in predicting Ca^2+^ transients and no Ca^2+^ transients (**N**, training cell number F_5,942_=164.6, p<0.01; testing cell source F_2,942_=18.3, p<0.01; training cell number × testing cell source interaction F_10,942_=0.3, p=0.98). When 20 cells were included in the machine learning training model, the macro F1 scores in predicting Ca^2+^ transients and no Ca^2+^ transients (**O**, F_2,157_=9.0, p<0.01) in testing cells from different regions in different mice reduced. **P,** Legends showing the color-coded abbreviations for 4 experimental groups. Data were analyzed by two-way ANOVA (**B**, **C**, **F**, **G**, **J**, **K**, **N**) or one-way ANOVA (**D**, **E**, **H**, **I**, **L**, **M**, **O**), followed by Bonferroni *post hoc* test. *, p<0.05; **, p<0.01. 40 testing cells in each group.

## 3. RESULTS

### 3.1. Comparison of the 4 machine learning models

Our goal was to find an efficient machine learning model with high predictability in detecting the Ca^2+^ transients while minimizing computational demands. To achieve this, we compared the performance of two RNN variants, GRU and LSTM, alongside two Transformer models, the standard Transformer and local Transformer. We used a dataset of 203 cells with manually confirmed Ca^2+^ transient as “ground” truth (see more details in Methods) (**Fig. 6A**). Our data showed that the macro F1 score for predicting Ca^2+^ transients and no Ca^2+^ transients was higher with RNN (GRU, 0.948 ± 0.002; LSTM, 0.946±0.004; mean ± S.E.M., same bellow) compared to the standard Transformer (0.919 ± 0.005), which was partially improved by local Transformer (0.935 ± 0.005) (**Fig. 6H**).

Specifically, for Ca^2+^ transient prediction, the F1 score was higher with RNN (GRU, 0.900 ± 0.005; LSTM, 0.894±0.008) compared to the standard Transformer (0.841 ± 0.010), which was partially improved by local Transformer (0.873 ± 0.010) (**Fig. 6F**). In terms of Precision, both GRU (0.873 ± 0.008) and LSTM (0.860 ± 0.010) models outperformed the standard Transformer (0.771 ± 0.015) and the local Transformer (0.835 ± 0.013) (**Fig. 6B**). However, the Recall scores across 4 machine learning models were similar (GRU: 0.926 ± 0.008; LSTM: 0.931 ± 0.010; standard Transformer: 0.929 ± 0.009; local Transformer: 0.916 ± 0.010) (**Fig. 6D**), indicating that the F1 score differences were mainly driven by Precision.

For predicting non-Ca^2+^ transients, the precision, recall, and F1 scores were all near perfect (>0.990), due to the consistent nature of non-Ca^2+^ transient signals. When analyzing F1 scores specifically for non-Ca^2+^ transient prediction, similar trends as the macro F1 scores were observed across the four machine learning models (**Fig. 6G**), with differences primarily driven by Recall (**Fig. 6E**) rather than Precision (**Fig. 6C**).

Compared to Transformer models, RNN models exhibited higher Precision in detecting Ca^2+^ transients and better Recall for non-Ca^2+^ transients, both attributable to RNN’s ability to minimize misclassification of the non-Ca^2+^ transient signal as Ca^2+^ transient signals. The consistent good Recall for Ca^2+^ transients and Precision values for non Ca^2+^ transients across all machine learning models indicates that they have a low likelihood of misclassifying Ca^2+^ transients as non-Ca^2+^ transients. The primary challenge in identifying Ca^2+^ transients lies in reducing errors where non-Ca^2+^ transients are mistakenly classified as transients. RNN models outperformed Transformer in addressing this issue, which was partially improved when using the local Transformer. Given its lower variability and faster processing time, GRU is recommended over LSTM. Therefore, all subsequent analyses were conducted using the GRU model.

### 3.2. Optimization of the cell number of the training data set

To determine the optimal number of cells to be used as the training dataset, we tested the predictability of GRU model trained by varying numbers of cells, ranging from 1 to 20. When the testing data sets were randomly sampled from the same session stage (i.e., the first 15 min or the last 15 min) in the same 1hr daily session on either Day 1 or Day 5, we observed a significant improvement in the macro F1 score when 5 to 20 cells were sampled, compared to GRU models trained with only 1 or 2 cells. However, no differences were observed among models trained with 5 to 20 cells (macro F1 score: 1 cell, 0.817 ± 0.014; 2 cells, 0.885 ± 0.008; 5 cells, 0.926 ± 0.005; 10 cells, 0.945 ± 0.004; 15 cells, 0.947 ± 0.004; 20 cells, 0.946 ± 0.004) (**Fig. 7H**). This cell number-dependent predictability is primarily attributable to the model’s predictability of the Ca^2+^ transient indicated by its Precision (**Fig. 7B**), Recall (**Fig. 7D**), and F1 scores (**Fig. 7F**) for Ca^2+^ transients, as the Precision (**Fig. 7C**), Recall (**Fig. 7E**) and F1 scores (**Fig. 7G**) for non-Ca^2+^ transient were all near perfect regardless of the cell number of the training data set. We further explored the effects of cell number when testing cells were randomly sampled from the different session stages within the same 1-hr daily session, the same session stage on a different daily session, and the different sessions stage on a different daily session of the same mouse (**Fig. 8A**). In all of these three cases, we found significant improvements of both Ca^2+^ transient prediction (**Fig. 8B, F, J, N**) and the non-Ca^2+^ transient predictions, though the latter were near perfect in most instances (**Fig. 8 C, G, K, O**), when 5 or more cells were sampled for training, compared to the models trained with only 1 or 2 cells. Finally, we extended further by randomly sampling the training data sets from different mice (**Fig. 9A**). We found significant improvements of Ca^2+^ transient prediction (**Fig. 9 B, F, J, N**) and the non-Ca^2+^ transient prediction, though the latter was near perfect in most instances (**Fig. 9 C, G, K, O**), when using 5 or more cells, regardless of whether the testing cells were from the same or different brain regions in mice distinct from those used for training. In conclusion, the number of cells used for training is crucial for accurately predicting Ca^2+^ transients. A minimum of 10 cells appears to be sufficient for achieving macro F1 scores above 0.900 when training the GRU model.

### 3.3. Sharing the machine learning model for predicting testing cells across different time windows in the same mouse

To assess how well a GRU model trained on data from a section of a 1-hr daily session can predict Ca^2+^ transients from any other recordings in the same mouse, we randomly sampled testing cells under four conditions: (1) the same session stage in the same daily session (smSS::smDay), (2) a different session stage in the same daily session (dfSS::smDay), (3) the same session stage in a different daily session (smSS::dfDay), and (4) a different session stage in a different daily session (dfSS::dfDay) (**Fig. 8A**). When 20 cells were sampled, the macro F1 scores were 0.946 ± 0.004 for smSS::smDay, 0.936 ± 0.006 for dfSS::smDay, 0.934 ± 0.005 for smSS::dfDay, and 0.927 ± 0.008 for dfSS::dfDay (**Fig. 8N**). Thus overall, the predictability remained high when testing cells were sampled from the same mouse used for GRU model training, though the macro F1 score for predicting both Ca^2+^ transients and non-Ca^2+^ transients was affected statistically, but mildly, by the source of testing cells (**Fig. 8 N, O**).

Specifically, for predicting Ca^2+^ transients, the source of testing cells affected the F1 scores (**Fig. 8 J, L**), particularly the Precision (**Fig. 8 B, D**) rather than the Recall (**Fig. 8 F, H**). When smSS::smDay was set as the gold standard with 20 cells as the benchmark, similar predictability was observed when testing cells were sampled from dfSS::smDay or smSS::dfDay. However, lower Precision and F1 score, but comparable Recall, were noted when testing cells were sampled from dfSS::dfDay.

For predicting non-Ca^2+^ transients, the source also affected the F1 scores (**Fig. 8 K, M**), primarily impacting the Recall (**Fig. 8 G, I**), while the Precision remained largely unaffected (**Fig. 8 C, E**). Similarly, as what was observed for predicting Ca^2+^ transients, we found the non-Ca^2+^ transient prediction was comparable when testing cells were sampled from smSS::smDay, dfSS::smDay or smSS::dfDay. Lower Recall and F1 score, but similar Precision, for non-Ca^2+^ transient prediction was detected when testing cells were sampled from dfSS::dfDay.

In conclusion, the GRU model can effectively predict Ca^2+^ transients in cells from the same mouse used for training. However, the risk of misclassifying non-Ca^2+^ transients as Ca^2+^ transients increases, though within a narrow range, when testing cells are sampled from a different session stage and a different day.

### 3.4. Sharing the machine learning model for predicting testing cells from different mice

To evaluate how the GRU model can predict the Ca^2+^ transients from a different mouse, either sharing the same brain region or switching to a different brain region, we used the smSS::smDay sampling strategy as the benchmark and added two more strategies in sampling testing cells: (1) the same brain region in a different mouse (denoted smRegion::dfMouse, and (2) a different brain region in a different mouse (denoted dfRegion::dfMouse). When 20 cells were sampled, the macro F1 scores were 0.946 ± 0.004 for smDay::smSS, 0.947 ± 0.003 for smRegion::dfMouse, and 0.927 ± 0.005 dfRegion::dfMouse. Thus, regardless of the brain region specificity, overall predictability remained high when testing cells were sampled from mice different from the one used for GRU training, although the macro F1 score for predicting both Ca^2+^ and non-Ca^2+^ transients was statistically reduced when the testing cells were sampled in dfRegion::dfMouse (**Fig. 9 N, O**).

Specifically, for predicting Ca^2+^ transients in different mice, the source of testing cells affected the F1 scores (**Fig. 9 J, L**) by impacting both the Precision (**Fig. 9 B, D**) and the Recall (**Fig. 9 F, H**). Lower Precision, Recall and F1 score were noted when testing cells were sampled from dfRegion::dfMouse, relative to either smSS::smDay or dfRegion::dfMouse.

For predicting non-Ca^2+^ transients, the source also affected the F1 scores (**Fig. 9 K, M**), by impacting both the Precision (**Fig. 8 C, E**) and the Recall (**Fig. 8 G, I**). Lower, although still near perfect, Precision, Recall and F1 score were noted when testing cells were sampled from dfRegion::dfMouse, relative to either smSS::smDay or dfRegion::dfMouse.

In conclusion, the GRU model can identify Ca^2+^ transients in cells from different mice with a macro F1 beyond 0.900. However, the risk of misclassifying non-Ca^2+^ transients as Ca^2+^ transients or Ca^2+^ transients as non-Ca^2+^ transients increases, though within a narrow range, when testing cells are sampled from a different brain region of a different mouse. There appears to be no significant impact on the F1 score when the testing cells are from the same region of a different mouse.

## 4. DISCUSSION

### 4.1. Selection of Machine Learning Model

In recent years, several studies using different machine learning methods have been developed utilizing some combination of the CNN, Attention-based architectures for cell identification and subsequent cell extraction^28–30^. However, they require a significant amount of labelled data as ground truth to achieve adequate predictive performance. This raises concerns in both time investment to train a single model as well as, due to the black box nature of these models, their ability to maintain performance given a substantial change in the input data (such as changes to different experimental animals, another brain region of interest, a different type of neuron).

In this study, we evaluated four machine learning models (GRU, LSTM, Transformer, and Local Transformer) for Ca^2+^ transient detection. Both predictability performance (i.e., precision, recall, and F1 scores) and time efficiency should be considered. **First**, GRU model is the most time-efficient, with a computation time of 287.1 second / epoch, slightly faster than LSTM (290.59 second / epoch) due to its simpler architecture and fewer parameters, making it computationally less expensive. GRU also exhibits strong overall performance in terms of precision, recall, and F1 scores, making it an excellent choice when considering both predictability and computation time. **Second**, LSTM takes marginally longer than GRU but performs slightly worse across multiple predictability metrics for Ca^2+^ transient detection. While very close to GRU in time efficiency, its more complex gating mechanism results in slightly longer computation times, without a significant performance advantage. **Third**, standard Transformer requires significantly more computation time at 708.8 second / epoch. Although transformers can be easily parallelized, their global self-attention mechanism and large parameter space demand much higher computational power. This increased time is not justified by any improvement in Ca^2+^ transient predictability, which actually suffers due to interruptions from distal frames when global attention is used in decision-making. **Fourth**, Local Transformer is the least time-efficient, taking 1744.4 second / epoch, almost 6x slower than GRU or LSTM, and 2.5x slower than the standard Transformer. This can be attributed to the increased complexity of managing localized attention windows and maintaining context between them. Additionally, it loses parallelization efficiency when processing the entire window. However, this increased computation time is compensated by improved predictability, as attention is focused more effectively on local frames, leading to better decision-making. **In conclusion**, RNNs outperform transformers in both time efficiency and predictability for Ca^2+^ transient detection. Between two RNN variants, GRU offers a better performance in both time efficiency and predictability, although LSTM remains a viable alternative. When the GRU performance is challenged by dataset size, computational resources, and the complexity of the transient dynamics, other models (e.g., LSTM) could be explored directly in the CalTrig.

### 4.2. Integrative Visual Exploration

The synchronization of Ca^2+^ imaging, cellular footprint, behavioral tracking, and Ca^2+^ transient trace windows, adjustable *via* a time bar, allows for a comprehensive and temporally aligned visualization of neuronal activity and behavior. This integration of multiple data streams provides several key benefits for neuroscience research. **First**, enhanced understanding of brain-behavior relationships: By synchronizing Ca^2+^ traces with behavioral data, researchers can directly observe how specific neuronal populations respond during behaviors or external stimuli. This real-time insight deepens our understanding of how brain activity drives behavior, such as the timing of Ca^2+^ transients relative to actions like lever presses. **Second**, technical flexibility and real-time adjustments: The adjustable time window enables researchers to zoom in on specific segments of data, closely examining how neural activity changes just before or after a behavioral event, which is critical for understanding causal relationships between stimuli and neural responses. **Third**, hypothesis proposing and testing: The integration of data streams facilitates hypothesis generation, which can be rapidly tested by determining the relationships between neural signals and behaviors. Researchers can observe whether certain Ca^2+^ transients precede or follow behavioral events, helping to refine hypotheses in real-time. **Fourth**, visualization of micro-network dynamics: Viewing neural footprints while tracking behavior enables the identification of micro-networks at the neuronal level, distinct from traditional brain region-level network, during specific tasks or stimuli. This is critical for understanding neuronal network dynamics in healthy and diseased states, such as in learning, memory, or addiction studies. **Fifth**, categorization of neurons: Visualizing Ca^2+^ transients allows for the clustering of neurons based on activity patterns, helping to uncover subpopulations with distinct response profiles. This helps researchers understand the heterogeneity and complexity of neural networks and their roles in both normal and pathological behaviors. **Sixth**, facilitating longitudinal studies: In long-term studies, where animals undergo repeated trials, integrative visualization tools allow researchers to track changes in both neuronal activity and behavior over time, aiding in the study of neuroplasticity and how neuronal responses evolve with experience. **Seventh**, improved data interpretation and collaboration: The ability to visualize multiple layers of data simultaneously aids in the interpretation of complex datasets. This holistic view helps researchers identify patterns and facilitates collaboration by making findings more accessible and easier to communicate across disciplines. **In conclusion**, integrative visual exploration enables deeper exploration of brain function, supports hypothesis generation about neuronal mechanisms, and opens new avenues for understanding neurological disorders and developing treatments.

### 4.3. Featured Value of CalTrig

The **<u>usability</u>** of the CalTrig tool is highlighted by its integrated visual experience, which provides users with a comprehensive view of Ca^2+^ images, neuron footprints, and Ca^2+^ transient traces in one unified interface. The tool features multi-layered toolboxes that cater to various analytical needs, allowing users to work efficiently at both the cellular and Ca^2+^ transient levels. Manual identification of Ca^2+^ transients is made efficient with features such as keyboard shortcuts (ASDF keys), hide/show traces, and unified scales for transient identification. This allows users to manually identify transients in approximately 1-2 minutes per 15-minute Ca^2+^ trace. Furthermore, manual identification can be assisted by auto-identification processes based on kinetic parameters or pre-established machine learning models. CalTrig is a self-looped tool, designed for continuous functional improvement, ensuring it remains adaptable and upgradable.

In terms of **<u>accessibility</u>**, CalTrig is a GUI-based open-source tool, making it readily shareable and open for further updates. It supports all CNMF or CNMF-E-processed imaging data and operates with limited computing demands, allowing for efficient performance even on less powerful systems. The GUI of CalTrig is designed to be user-friendly, ensuring that non-programmers can easily interact with it through intuitive buttons, menus, and forms. The tool presents a professional appearance, which enhances its suitability for presentations, collaborations, and publications. Real-time feedback is provided during parameter adjustments in Ca^2+^ transient identification, allowing users to fine-tune the process interactively. CalTrig can be downloaded as an independent application that runs in a Python environment, meaning users do not need additional software installations. Additionally, the tool is highly customizable, allowing users to tailor it to specific workflows and improving productivity for different use cases. Its expandability ensures that it can accommodate future updates and feature integrations.

Our research demonstrates that the GRU model provides high predictability when applied to testing cells from the same or different session stages, across various training days, brain regions, and mice. The GRU model has proven to be efficient in training, as shown in **Fig. 7**, even with a limited number of cells (Figs. **8****-10**), and the “ground truth” of Ca^2+^ transients can be established with relative ease, as detailed in the **Methods** and **Results** sections. This makes it highly feasible to extend testing to longer training sessions, such as the 6-hour sessions commonly used in studies on learning, memory, motivation, and addiction, or across broader time spans covering days, weeks, or months. The tool we developed here will be instrumental in evaluating the feasibility of *in vivo* Ca^2+^ recordings during extended sessions over prolonged recording periods.

The high predictability of the GRU model in detecting Ca^2+^ transients also indicates that the basic properties of Ca^2+^ transients, such as rise slope, amplitude, and signal-to-noise ratio, remained stable during 1-hour recording sessions, across five recording days, and across different brain regions in several mice. Notably, the neurons detected in two brain regions, M2 and PrL, are most likely pyramidal neurons, which may explain the model’s high level of expandability. A future direction for research could be to test the expandability of the model to different types of neurons, e.g., interneurons.

### 4.4. Limitations and Future Directions

There are several areas which the current study has not yet explored. For instance, we have not tested the trained machine learning model for detecting Ca^2+^ transients across different animal lines or across species, such as from mice to rats. The Ca^2+^ indicator used in this study is GCaMP8f, and it would be interesting to compare the dynamics of Ca^2+^ transients and the persistence of fluorescent intensity using other genetically encoded Ca^2+^ indicators (GECIs). Furthermore, the detection of Ca^2+^ transients is often not the end goal of data analysis. When conducting longitudinal recordings across different experimental groups, researchers may face challenges related to selecting appropriate time windows for analysis. Behavior-associated Ca^2+^ transients present a particularly valuable area for exploration, as they offer insights into neuronal activities linked to specific brain functions. Another area of interest is the association between neuronal footprints and activity in specific physiological or pathological conditions. This is a largely unexplored field, but it is intriguing to consider how the spatial distribution of neurons may influence brain function. Our team is currently developing relevant tools to integrate with CalTrig, which will further enhance our ability to understand the role of Ca^2+^ transients in brain output coding.

## ACKNOWLEDGEMENTS

This work was supported by NIH grants (R01AG072897, R01AA025784, and R01DA059548). We thank the support from High-Performance Computing and Storage at Indiana University. This work also received support from the Indiana Spinal Cord & Brain Injury Research Fund (Indiana State Department of Health); its contents are solely the responsibility of the providers and do not necessarily represent the official views of the Indiana State Department of Health. We acknowledge Dr. Denise Cai and her team for developing MiniAn as a powerful tool used to extract Ca^2+^ traces.

## AUTHOR CONTRIBUTION

Experimental design: MAL, YC, HF, CG, YYM. Data collection: MAL, YC. Data analysis: MAL, YC, AK, CG, YM. Results, Discussion: MAL, YC, HF, AK, CG, YYM. Manuscript preparation: MAL, YC, HF, AK, CG, YYM.

