## Supplementary Information and figures for "CalTrig: A GUI-based Machine Learning Approach for Decoding Neuronal Calcium Transients in Freely Moving Rodents"

***Corresponding author**: Dr. Yao-Ying Ma

### Signal-to-Noise Estimation

While verifying transient events, occasional spikes were detected due to background fluctuations rather than specific cellular activity, and in rare cases, noise was misinterpreted as calcium signals. Contemporary methods often use z-scores to estimate noise, but this global noise estimation lacks robustness when noise fluctuates over time, potentially leading to the rejection of valid transients during low-noise periods.

To address this concern, we apply the Savitzky–Golay (SavGol) filter (**Fig. S6**), which smooths noisy data while preserving important features such as peaks and edges (Steinier, Termonia et al. 1972, Dai, Selesnick et al. 2017). This filter fits successive polynomial functions to adjacent data points, making it ideal for time-series data where preserving peak shape is crucial. In our pilot studies, where we compared the original ΔF/F with the smoothed ΔF/F using CalTrig's data exploration tools, the Ca2+ transients were well-preserved. Therefore, the SavGol filter was chosen as the primary smoothing filter.

Noise estimation for each image frame is calculated by the difference between the original and SavGol-filtered ΔF/F:

$$Sig_{noise}(t) \approx \left| Sig_{\frac{\Delta F}{F}} - savgol\left(Sig_{\frac{\Delta F}{F}}\right) \right|$$

To further refine the noise estimation, we apply a rolling window function, preventing small value overlaps between the smoothed and non-smoothed signals. The user can choose smoothing types (average, median, max) and adjust the rolling window size (20 frames is a recommended starting point). SNR is calculated by dividing the smoothed signal by the estimated noise, with a minimum cap (e.g., 0.1) to avoid ineffective or exaggerated values. The resulting SNR effectively mirrors the ΔF/F, proportionally accentuated or diminished by the noise.

The parameters for the SavGol filter and noise smoothing can be adjusted under the "SavGol" and "Noise" subtabs in the Trace toolbox window. Visualization options for the SavGol-filtered ΔF/F, noise, and SNR are available under the "Params" subtab for easy analysis in the $Ca^{2+}$ trace window.

### SUPPLEMENTARY FIGURES

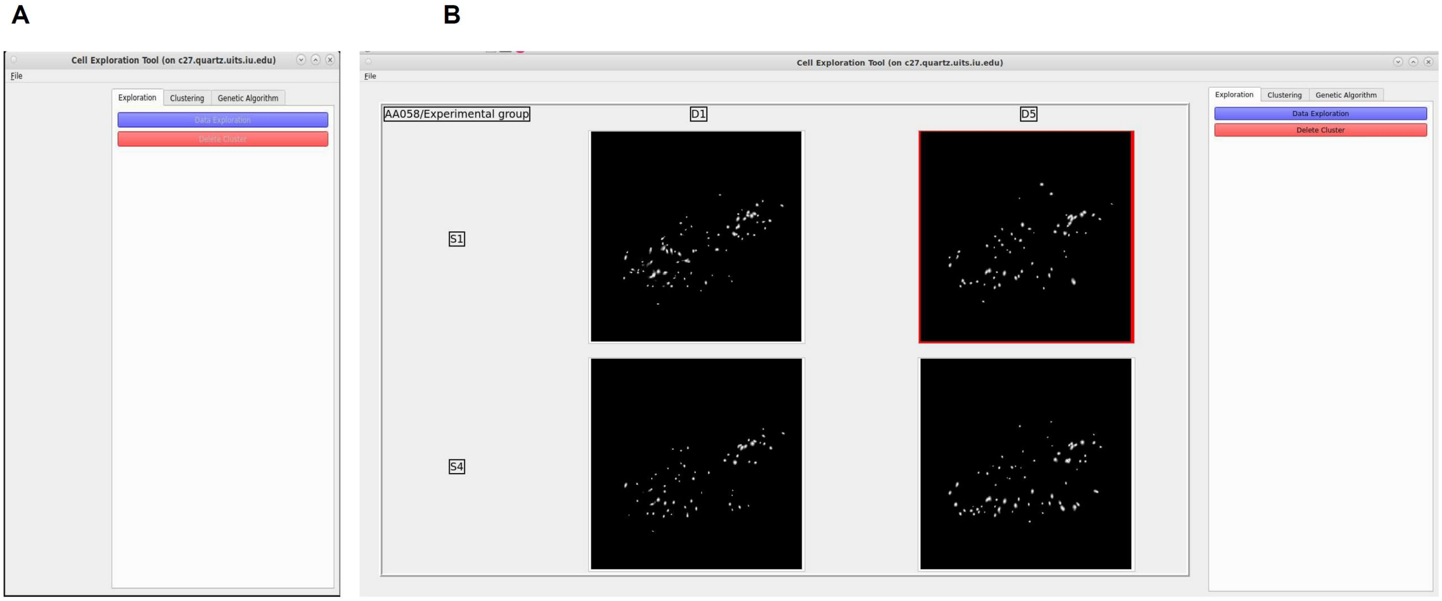


#### Figure S1. Data loading

**A**, The front page of CalTrig serves as a hub for loading data

**B**, Each set of data can be visualized in the window by their cell footprint and are arranged in a grid related to the experiment design details, such as animal ID, day, session stage

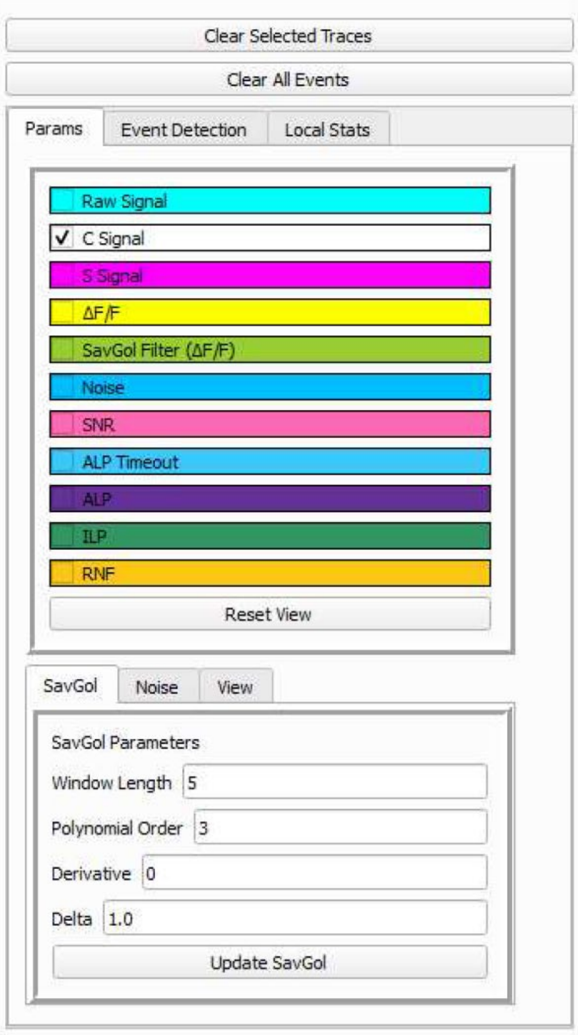


**Figure S2. Parameter list**

In upper panel of the Trace Toolbox, raw signal, C signal, S signal, ΔF/F, Noise, Signal to noise ratio (SNR), as well as other behavioral readouts (e.g., active lever press, ALP; ALP during timeout; inactive lever press; and reinforcement, RNF) are listed as items to be selected to show in the $Ca^{2+}$ Trace window. The parameters of Savitzky–Golay filter (SavGol), used to estimate the SNR, are listed in the lower panel.

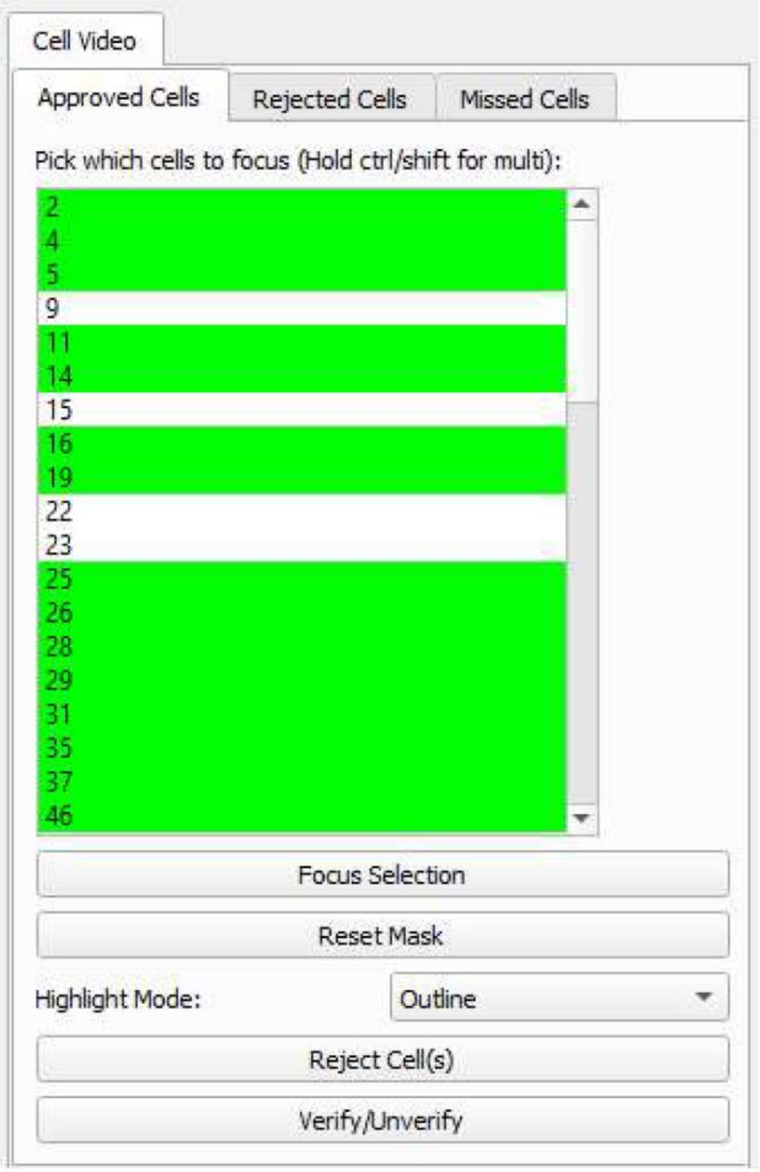


**Figure S3. Cell verification**

CalTrig allows user to inspect individual cells detected by CalV2N by interactive exploration of the $Ca^{2+}$ imaging window and $Ca^{2+}$ trace window. A cell becomes verified after a visual confirmation of its footprint, $Ca^{2+}$ Image, and $Ca^{2+}$ trace, ensuring identifiable $Ca^{2+}$ transients are present. The verified cells are marked by yellow highlight.

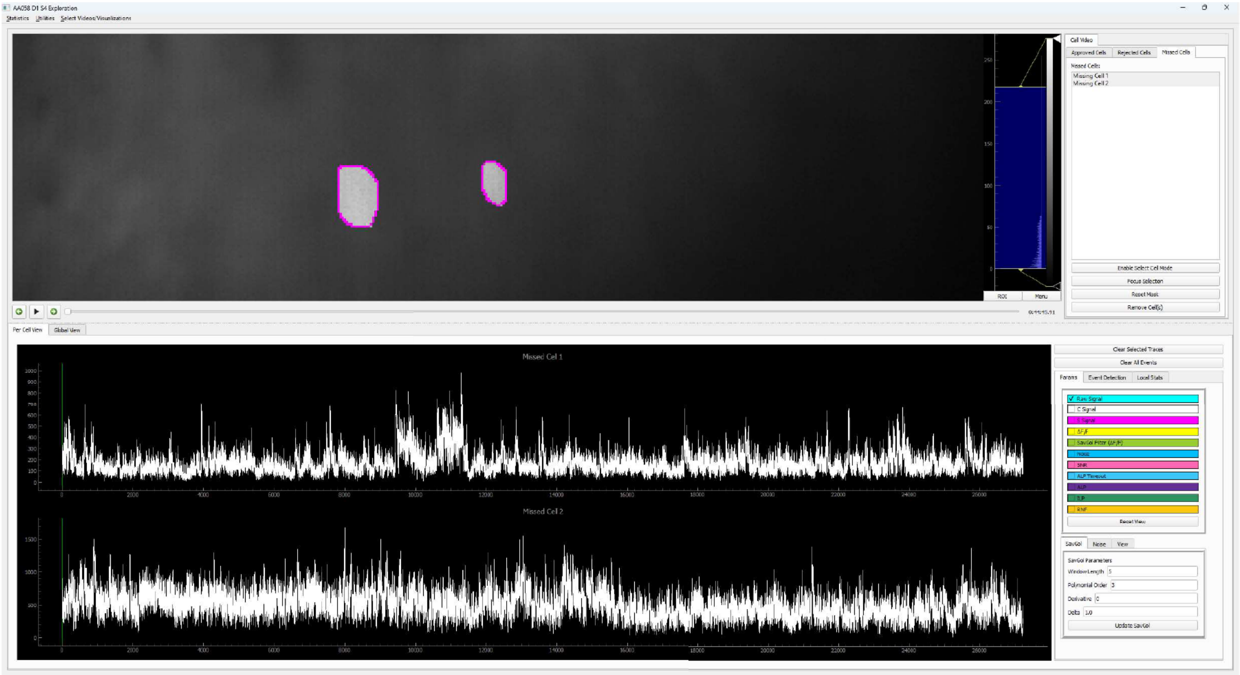


**Figure S4. Add missing cell**

While reviewing the $Ca^{2+}$ imaging video, one may notice a potentially missed cell by CalV2N. CalTrig allows user to manually draw the contour of the suspected cell in the $Ca^{2+}$ imaging window, creating its footprint. The corresponding temporal trace of signal intensity for the selected area is then calculated by averaging pixel intensities and displayed in the $Ca^{2+}$ trace window. If the $Ca^{2+}$ traces meet the criteria for accepted cells, the manually identified cell can be added to the list of missing cells.

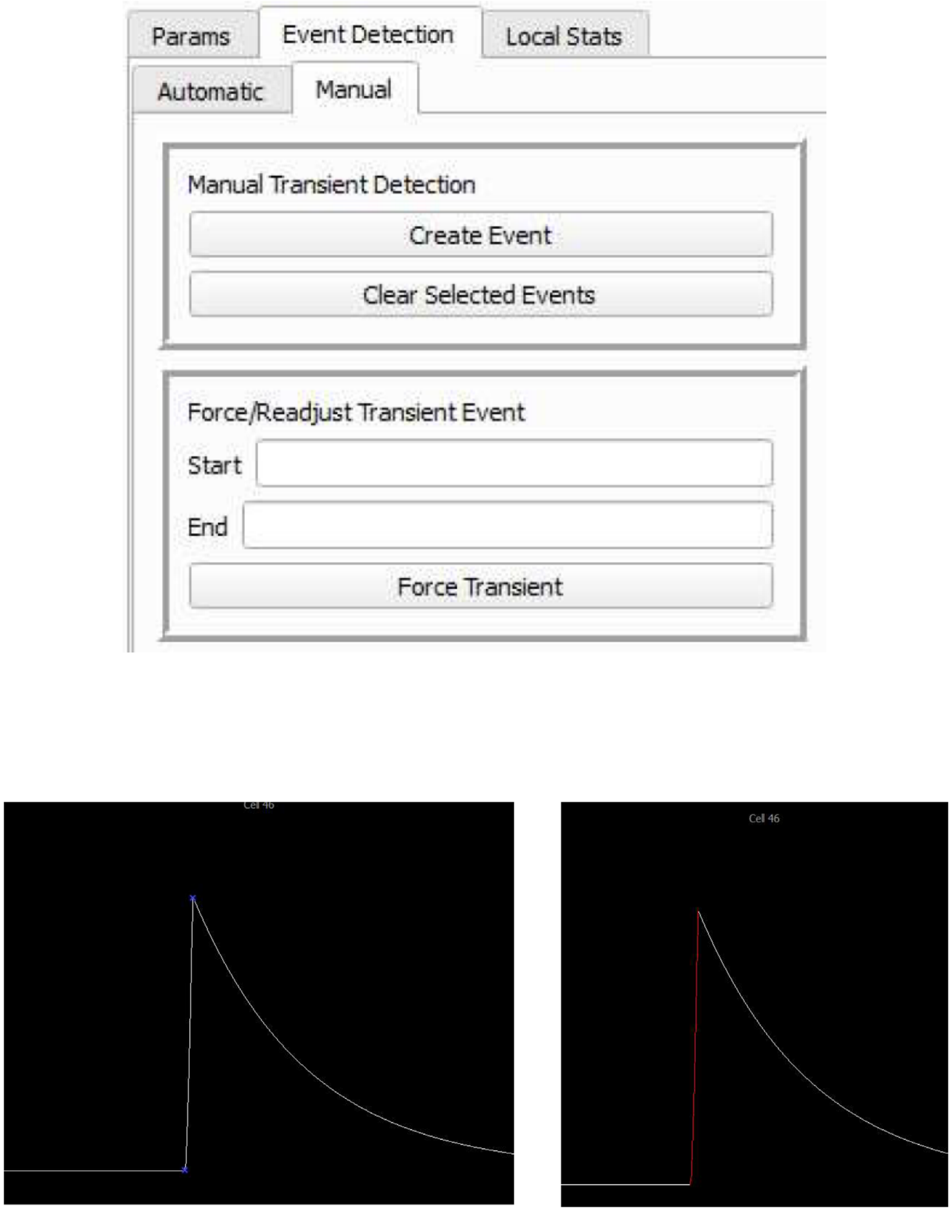


**Figure S5. Manual identification of $Ca^{2+}$ transients**

$Ca^{2+}$ transients can be directly identified by manually selecting the start and end points of the rising section of $Ca^{2+}$ transients.

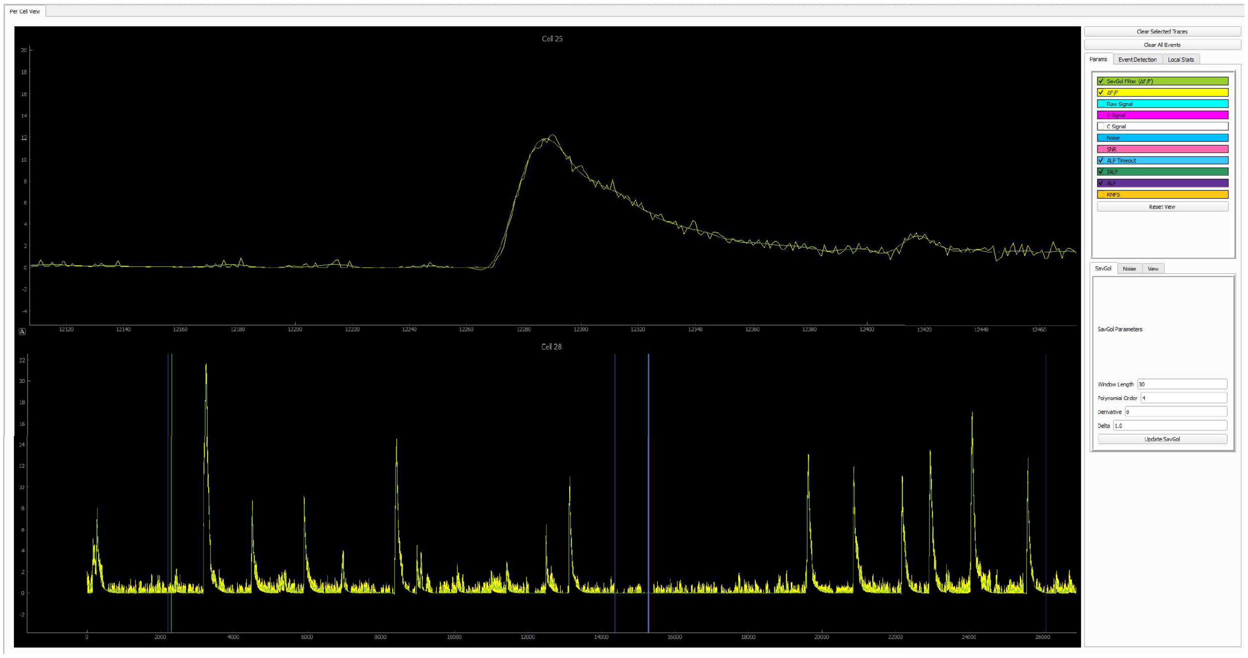


### Figure S6. Savitzky–Golay filter

This sets the minimal Signal-to-Noise ratio (SNR). Using the Savitzky–Golay filter [12,13] to smooth ΔF/F signals, noise is calculated as the difference between the original and filtered ΔF/F, further smoothed by a rolling window strategy. Then SNR is computed by dividing the smoothed ΔF/F by the estimated noise.

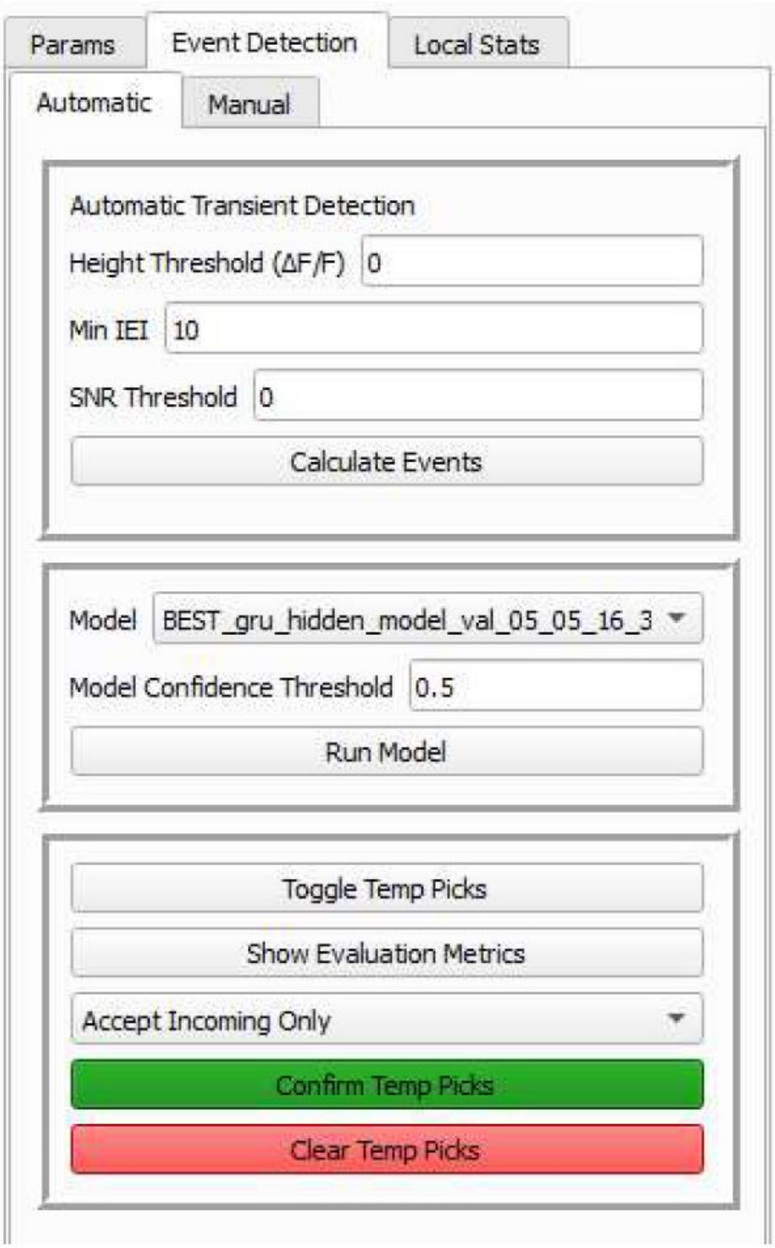


**Figure S7. Automatic detection**

To enhance detection accuracy, the parameter-based auto detection algorithm starts by including all C peaks in a candidate pool, then filtering down to a valid subset based on three pre-defined parameters: "Peak Threshold (ΔF/F)", "Interval Threshold", and "SNR Threshold".

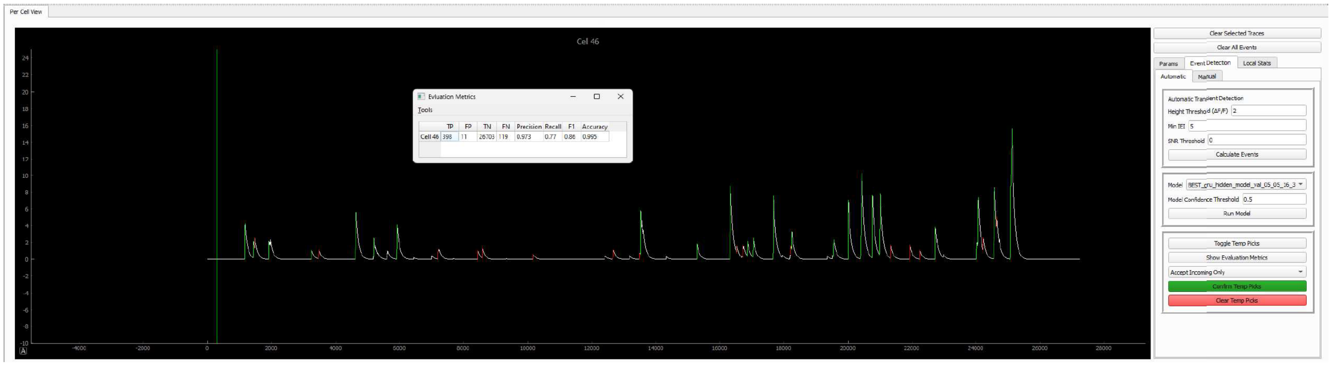


**Figure S8. Machine learning evaluation**

We used Precision, Recall, F1 and macro F1 as key metrics to evaluate the performance of machine learning model in predicting $Ca^{2+}$ transients *vs*. non-$Ca^{2+}$ transients.

A

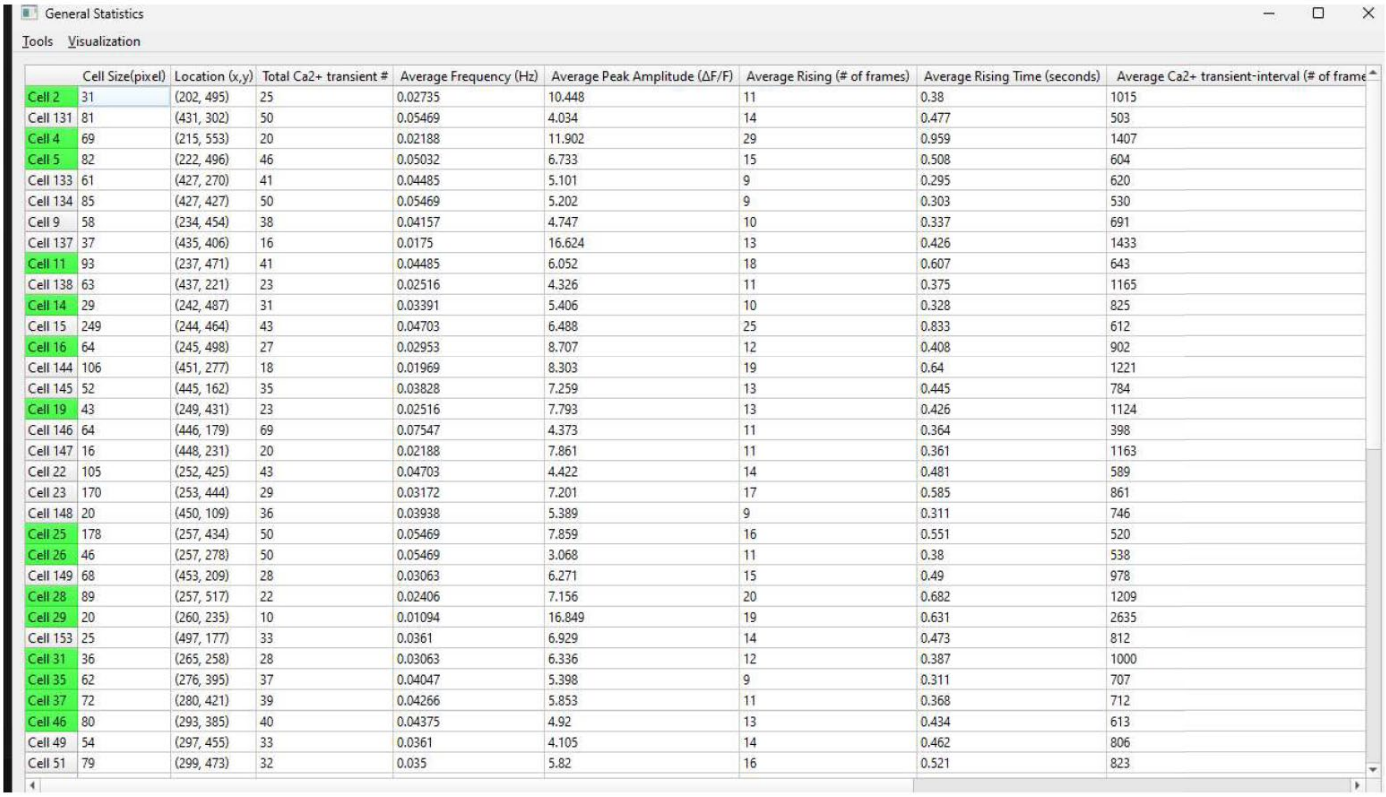

General Statistics

Tools Visualization

| | Cell Size(pixel) | Location (x,y) | Total Ca2+ transient # | Average Frequency (Hz) | Average Peak Amplitude (ΔF/F) | Average Rising (# of frames) | Average Rising Time (seconds) | Average Ca2+ transient-interval (# of frame |
|---|---|---|---|---|---|---|---|---|
| Cell 2 | 31 | (202, 495) | 25 | 0.02735 | 10.448 | 11 | 0.38 | 1015 |
| Cell 131 | 81 | (431, 302) | 50 | 0.05469 | 4.034 | 14 | 0.477 | 503 |
| Cell 4 | 69 | (215, 553) | 20 | 0.02188 | 11.902 | 29 | 0.959 | 1407 |
| Cell 5 | 82 | (222, 496) | 46 | 0.05032 | 6.733 | 15 | 0.508 | 604 |
| Cell 133 | 61 | (427, 270) | 41 | 0.04485 | 5.101 | 9 | 0.295 | 620 |
| Cell 134 | 85 | (427, 427) | 50 | 0.05469 | 5.202 | 9 | 0.303 | 530 |
| Cell 9 | 58 | (234, 454) | 38 | 0.04157 | 4.747 | 10 | 0.337 | 691 |
| Cell 137 | 37 | (435, 406) | 16 | 0.0175 | 16.624 | 13 | 0.426 | 1433 |
| Cell 11 | 93 | (237, 471) | 41 | 0.04485 | 6.052 | 18 | 0.607 | 643 |
| Cell 138 | 63 | (437, 221) | 23 | 0.02516 | 4.326 | 11 | 0.375 | 1165 |
| Cell 14 | 29 | (242, 487) | 31 | 0.03391 | 5.406 | 10 | 0.328 | 825 |
| Cell 15 | 249 | (244, 464) | 43 | 0.04703 | 6.488 | 25 | 0.833 | 612 |
| Cell 16 | 64 | (245, 498) | 27 | 0.02953 | 8.707 | 12 | 0.408 | 902 |
| Cell 144 | 106 | (451, 277) | 18 | 0.01969 | 8.303 | 19 | 0.64 | 1221 |
| Cell 145 | 52 | (445, 162) | 35 | 0.03828 | 7.259 | 13 | 0.445 | 784 |
| Cell 19 | 43 | (249, 431) | 23 | 0.02516 | 7.793 | 13 | 0.426 | 1124 |
| Cell 146 | 64 | (446, 179) | 69 | 0.07547 | 4.373 | 11 | 0.364 | 398 |
| Cell 147 | 16 | (448, 231) | 20 | 0.02188 | 7.861 | 11 | 0.361 | 1163 |
| Cell 22 | 105 | (252, 425) | 43 | 0.04703 | 4.422 | 14 | 0.481 | 589 |
| Cell 23 | 170 | (253, 444) | 29 | 0.03172 | 7.201 | 17 | 0.585 | 861 |
| Cell 148 | 20 | (450, 109) | 36 | 0.03938 | 5.389 | 9 | 0.311 | 746 |
| Cell 25 | 178 | (257, 434) | 50 | 0.05469 | 7.859 | 16 | 0.551 | 520 |
| Cell 26 | 46 | (257, 278) | 50 | 0.05469 | 3.068 | 11 | 0.38 | 538 |
| Cell 149 | 68 | (453, 209) | 28 | 0.03063 | 6.271 | 15 | 0.49 | 978 |
| Cell 28 | 89 | (257, 517) | 22 | 0.02406 | 7.156 | 20 | 0.682 | 1209 |
| Cell 29 | 20 | (260, 235) | 10 | 0.01094 | 16.849 | 19 | 0.631 | 2635 |
| Cell 153 | 25 | (497, 177) | 33 | 0.0361 | 6.929 | 14 | 0.473 | 812 |
| Cell 31 | 36 | (265, 258) | 28 | 0.03063 | 6.336 | 12 | 0.387 | 1000 |
| Cell 35 | 62 | (276, 395) | 37 | 0.04047 | 5.398 | 9 | 0.311 | 707 |
| Cell 37 | 72 | (280, 421) | 39 | 0.04266 | 5.853 | 11 | 0.368 | 712 |
| Cell 46 | 80 | (293, 385) | 40 | 0.04375 | 4.92 | 13 | 0.434 | 613 |
| Cell 49 | 54 | (297, 455) | 33 | 0.0361 | 4.105 | 14 | 0.462 | 806 |
| Cell 51 | 79 | (299, 473) | 32 | 0.035 | 5.82 | 16 | 0.521 | 823 |

B

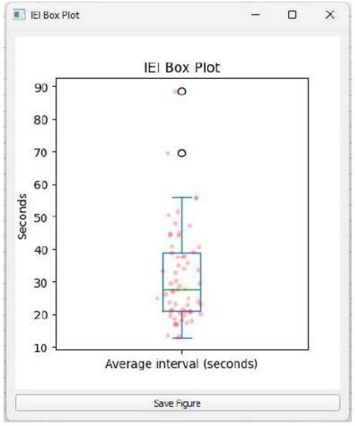


C

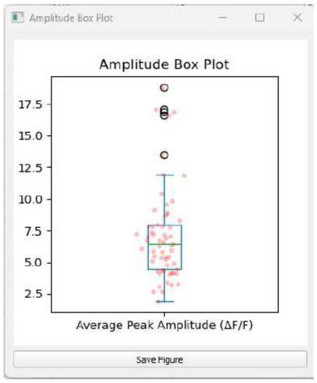

**Figure S9. Animal-wide data export**

**A**, A data table is generated, with each row representing one of the CalS2N-detected cells and each column providing specific readout as a general description of that cell. **B**, **C**, Average Peak Amplitude (**B**) and inter-transient interval (ITI, **C**), can be directly generated and saved as editable SVG files for publication purposes.

A

| | Rising-Start(frames) | Rising-Stop(frames) | Total # of Rising Frames | Rising-Start(seconds) | Rising-Stop(seconds) | Total # of Rising Frames (seconds) | Interval with Previous Transient (frames) | Interval with Previous Transient (seconds) | Peak Amplitude (ΔF/F) | Total Amplitude (ΔF/F) |
|---|---|---|---|---|---|---|---|---|---|---|
| Transient 1 | 1164 | 1174 | 10 | 2724.956 | 2725.291 | 0.335 | N/A | N/A | 6.986 | 39.683 |
| Transient 2 | 1423 | 1432 | 9 | 2733.65 | 2733.954 | 0.304 | 259 | 8.66 | 2.578 | 16.618 |
| Transient 3 | 1471 | 1480 | 9 | 2735.262 | 2735.564 | 0.302 | 48 | 1.575 | 2.71 | 23.591 |
| Transient 4 | 1910 | 1922 | 12 | 2750.002 | 2750.405 | 0.403 | 439 | 14.703 | 2.916 | 25.353 |
| Transient 5 | 3242 | 3249 | 7 | 2794.723 | 2794.959 | 0.236 | 1332 | 44.687 | 2.532 | 10.212 |
| Transient 6 | 3474 | 3479 | 5 | 2802.51 | 2802.679 | 0.169 | 232 | 7.753 | 2.09 | 7.118 |
| Transient 7 | 4613 | 4637 | 24 | 2840.754 | 2841.558 | 0.804 | 1139 | 38.21 | 7.726 | 79.0 |
| Transient 8 | 5184 | 5192 | 8 | 2859.923 | 2860.191 | 0.268 | 571 | 19.137 | 4.901 | 23.036 |
| Transient 9 | 5907 | 5920 | 13 | 2884.199 | 2884.633 | 0.434 | 723 | 24.242 | 5.024 | 44.174 |
| Transient 10 | 7197 | 7205 | 8 | 2927.507 | 2927.776 | 0.269 | 1290 | 43.276 | 2.066 | 11.188 |
| Transient 11 | 8430 | 8435 | 5 | 2968.904 | 2969.071 | 0.167 | 1233 | 41.362 | 2.231 | 8.799 |
| Transient 12 | 8578 | 8583 | 5 | 2973.872 | 2974.042 | 0.17 | 148 | 4.934 | 3.128 | 9.572 |
| Transient 13 | 10161 | 10166 | 5 | 3027.021 | 3027.189 | 0.168 | 1583 | 53.115 | 1.549 | 4.962 |
| Transient 14 | 12657 | 12673 | 16 | 3110.824 | 3111.358 | 0.534 | 2496 | 83.771 | 1.646 | 15.674 |
| Transient 15 | 13488 | 13495 | 7 | 3138.722 | 3138.955 | 0.233 | 831 | 27.867 | 2.078 | 8.856 |
| Transient 16 | 13511 | 13526 | 15 | 3139.494 | 3139.997 | 0.503 | 23 | 0.738 | 7.375 | 70.288 |
| Transient 17 | 15283 | 15288 | 5 | 3198.986 | 3199.154 | 0.168 | 1772 | 59.459 | 3.114 | 11.304 |
| Transient 18 | 16316 | 16329 | 13 | 3233.667 | 3234.105 | 0.438 | 1033 | 34.648 | 10.677 | 85.5 |
| Transient 19 | 16527 | 16532 | 5 | 3240.752 | 3240.92 | 0.168 | 211 | 7.051 | 2.157 | 10.26 |
| Transient 20 | 16715 | 16722 | 7 | 3247.065 | 3247.3 | 0.235 | 188 | 6.277 | 1.777 | 9.111 |
| Transient 21 | 16858 | 16865 | 7 | 3251.868 | 3252.1 | 0.232 | 143 | 4.77 | 2.805 | 14.541 |
| Transient 22 | 17038 | 17045 | 7 | 3257.908 | 3258.143 | 0.235 | 180 | 6.009 | 3.519 | 17.107 |
| Transient 23 | 17659 | 17673 | 14 | 3278.759 | 3279.228 | 0.469 | 621 | 20.816 | 9.798 | 80.785 |
| Transient 24 | 18212 | 18218 | 6 | 3297.324 | 3297.526 | 0.202 | 553 | 18.531 | 2.695 | 13.749 |
| Transient 25 | 18241 | 18249 | 8 | 3298.298 | 3298.567 | 0.269 | 29 | 0.94 | 4.575 | 30.86 |
| Transient 26 | 19543 | 19561 | 18 | 3342.01 | 3342.616 | 0.606 | 1302 | 43.679 | 3.645 | 38.428 |
| Transient 27 | 20001 | 20013 | 12 | 3357.388 | 3357.791 | 0.403 | 458 | 15.342 | 8.802 | 68.818 |
| Transient 28 | 20388 | 20433 | 45 | 3370.382 | 3371.892 | 1.51 | 387 | 12.961 | 10.855 | 229.565 |
| Transient 29 | 20753 | 20766 | 13 | 3382.635 | 3383.073 | 0.438 | 365 | 12.22 | 8.661 | 64.859 |
| Transient 30 | 20978 | 21010 | 32 | 3390.191 | 3391.264 | 1.073 | 225 | 7.523 | 7.092 | 115.577 |
| Transient 31 | 21323 | 21330 | 7 | 3401.772 | 3402.007 | 0.235 | 345 | 11.544 | 2.8 | 13.589 |
| Transient 32 | 21925 | 21933 | 8 | 3421.986 | 3422.255 | 0.269 | 602 | 20.179 | 2.562 | 15.65 |
| Transient 33 | 22239 | 22245 | 6 | 3432.529 | 3432.729 | 0.2 | 314 | 10.508 | 1.862 | 10.296 |
| Transient 34 | 22715 | 22725 | 10 | 3448.507 | 3448.843 | 0.336 | 476 | 15.947 | 5.58 | 37.09 |
| Transient 35 | 24010 | 24031 | 21 | 3491.986 | 3492.691 | 0.705 | 1295 | 43.445 | 3.634 | 47.714 |
| Transient 36 | 24057 | 24069 | 12 | 3493.564 | 3493.967 | 0.403 | 47 | 1.544 | 11.194 | 80.885 |
| Transient 37 | 24220 | 24226 | 6 | 3499.038 | 3499.237 | 0.199 | 163 | 5.44 | 4.02 | 20.336 |
| Transient 38 | 24549 | 24577 | 28 | 3510.082 | 3511.022 | 0.94 | 329 | 11.013 | 10.244 | 153.565 |
| Transient 39 | 24639 | 24649 | 10 | 3513.104 | 3513.439 | 0.335 | 90 | 2.988 | 6.607 | 60.13 |
| Transient 40 | 25062 | 25121 | 59 | 3527.306 | 3529.286 | 1.98 | 423 | 14.166 | 17.947 | 609.054 |

B

Statistics for Cell 46

Tools Visualization

| | Rising-Start(frames) | Rising-Stop(frames) | Total # of Rising Frames | Rising-Start(seconds) | Rising-Stop(seconds) | Total # of Rising Frames (seconds) | Interval with Previous Transient (frames) | Interval with Previous Transient (seconds) | Peak Amplitude (ΔF/F) | Total Amplitude (ΔF/F) |
|---|---|---|---|---|---|---|---|---|---|---|
| Transient 13 | 10161 | 10166 | 5 | 3027.021 | 3027.189 | 0.168 | 1583 | 53.115 | 1.549 | 4.962 |
| Transient 6 | 3474 | 3479 | 5 | 2802.51 | 2802.679 | 0.169 | 232 | 7.753 | 2.09 | 7.118 |
| Transient 11 | 8430 | 8435 | 5 | 2968.904 | 2969.071 | 0.167 | 1233 | 41.362 | 2.231 | 8.799 |
| Transient 15 | 13488 | 13495 | 7 | 3138.722 | 3138.955 | 0.233 | 831 | 27.867 | 2.078 | 8.856 |
| Transient 20 | 16715 | 16722 | 7 | 3247.065 | 3247.3 | 0.235 | 188 | 6.277 | 1.777 | 9.111 |
| Transient 12 | 8578 | 8583 | 5 | 2973.872 | 2974.042 | 0.17 | 148 | 4.934 | 3.128 | 9.572 |
| Transient 5 | 3242 | 3249 | 7 | 2794.723 | 2794.959 | 0.236 | 1332 | 44.687 | 2.532 | 10.212 |
| Transient 19 | 16527 | 16532 | 5 | 3240.752 | 3240.92 | 0.168 | 211 | 7.051 | 2.157 | 10.26 |
| Transient 33 | 22239 | 22245 | 6 | 3432.529 | 3432.729 | 0.2 | 314 | 10.508 | 1.862 | 10.296 |
| Transient 10 | 7197 | 7205 | 8 | 2927.507 | 2927.776 | 0.269 | 1290 | 43.276 | 2.066 | 11.188 |
| Transient 17 | 15283 | 15288 | 5 | 3198.986 | 3199.154 | 0.168 | 1772 | 59.459 | 3.114 | 11.304 |
| Transient 31 | 21323 | 21330 | 7 | 3401.772 | 3402.007 | 0.235 | 345 | 11.544 | 2.8 | 13.589 |
| Transient 24 | 18212 | 18218 | 6 | 3297.324 | 3297.526 | 0.202 | 553 | 18.531 | 2.695 | 13.749 |
| Transient 21 | 16858 | 16865 | 7 | 3251.868 | 3252.1 | 0.232 | 143 | 4.77 | 2.805 | 14.541 |
| Transient 32 | 21925 | 21933 | 8 | 3421.986 | 3422.255 | 0.269 | 602 | 20.179 | 2.562 | 15.65 |
| Transient 14 | 12657 | 12673 | 16 | 3110.824 | 3111.358 | 0.534 | 2496 | 83.771 | 1.646 | 15.674 |
| Transient 2 | 1423 | 1432 | 9 | 2733.65 | 2733.954 | 0.304 | 259 | 8.66 | 2.578 | 16.618 |
| Transient 22 | 17038 | 17045 | 7 | 3257.908 | 3258.143 | 0.235 | 180 | 6.009 | 3.519 | 17.107 |
| Transient 37 | 24220 | 24226 | 6 | 3499.038 | 3499.237 | 0.199 | 163 | 5.44 | 4.02 | 20.336 |
| Transient 8 | 5184 | 5192 | 8 | 2859.923 | 2860.191 | 0.268 | 571 | 19.137 | 4.901 | 23.036 |
| Transient 3 | 1471 | 1480 | 9 | 2735.262 | 2735.564 | 0.302 | 48 | 1.575 | 2.71 | 23.591 |
| Transient 4 | 1910 | 1922 | 12 | 2750.002 | 2750.405 | 0.403 | 439 | 14.703 | 2.916 | 25.353 |
| Transient 25 | 18241 | 18249 | 8 | 3298.298 | 3298.567 | 0.269 | 29 | 0.94 | 4.575 | 30.86 |
| Transient 34 | 22715 | 22725 | 10 | 3448.507 | 3448.843 | 0.336 | 476 | 15.947 | 5.58 | 37.09 |
| Transient 26 | 19543 | 19561 | 18 | 3342.01 | 3342.616 | 0.606 | 1302 | 43.679 | 3.645 | 38.428 |
| Transient 1 | 1164 | 1174 | 10 | 2724.956 | 2725.291 | 0.335 | N/A | N/A | 6.986 | 39.683 |
| Transient 9 | 5907 | 5920 | 13 | 2884.199 | 2884.633 | 0.434 | 723 | 24.242 | 5.024 | 44.174 |
| Transient 35 | 24010 | 24031 | 21 | 3491.986 | 3492.691 | 0.705 | 1295 | 43.445 | 3.634 | 47.714 |
| Transient 39 | 24639 | 24649 | 10 | 3513.104 | 3513.439 | 0.335 | 90 | 2.988 | 6.607 | 60.13 |
| Transient 29 | 20753 | 20766 | 13 | 3382.635 | 3383.073 | 0.438 | 365 | 12.22 | 8.661 | 64.859 |
| Transient 27 | 20001 | 20013 | 12 | 3357.388 | 3357.791 | 0.403 | 458 | 15.342 | 8.802 | 68.818 |
| Transient 16 | 13511 | 13526 | 15 | 3139.494 | 3139.997 | 0.503 | 23 | 0.738 | 7.375 | 70.288 |
| Transient 7 | 4613 | 4637 | 24 | 2840.754 | 2841.558 | 0.804 | 1139 | 38.21 | 7.726 | 79.0 |
| Transient 23 | 17659 | 17673 | 14 | 3278.759 | 3279.228 | 0.469 | 621 | 20.816 | 9.798 | 80.785 |
| Transient 36 | 24057 | 24069 | 12 | 3493.564 | 3493.967 | 0.403 | 47 | 1.544 | 11.194 | 80.885 |
| Transient 18 | 16316 | 16329 | 13 | 3233.667 | 3234.105 | 0.438 | 1033 | 34.648 | 10.677 | 85.5 |
| Transient 30 | 20978 | 21010 | 32 | 3390.191 | 3391.264 | 1.073 | 225 | 7.523 | 7.092 | 115.577 |
| Transient 38 | 24549 | 24577 | 28 | 3510.082 | 3511.022 | 0.94 | 329 | 11.013 | 10.244 | 153.565 |
| Transient 28 | 20388 | 20433 | 45 | 3370.382 | 3371.892 | 1.51 | 387 | 12.961 | 10.855 | 229.565 |
| Transient 40 | 25062 | 25121 | 59 | 3527.306 | 3529.286 | 1.98 | 423 | 14.166 | 17.947 | 609.054 |

C

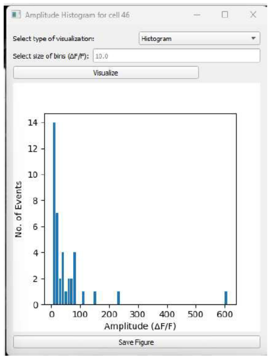


D

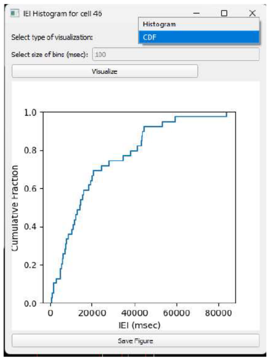

**Figure S10. Cell-wide data export**

**A**, A data table is generated for a selected cell, with the top row displaying column titles and the subsequent rows representing individual $Ca^{2+}$ transients detected by CalTrig, one transient per row.

**B**, The column title (e.g., Total Amplitude) can be selected to sort the data based on the information in the selected column.

**C**, **D**, Figures such as Amplitude Distribution (**C**) and ITI Frequency Histograms (**D**) can be created directly from this data and saved as editable SVG files for publication.
